# Molecular mechanisms for activation of the 26S proteasome

**DOI:** 10.1101/2023.05.09.540094

**Authors:** Donghoon Lee, Yanan Zhu, Louis Colson, Xiaorong Wang, Siyi Chen, Emre Tkacik, Lan Huang, Qi Ouyang, Alfred L. Goldberg, Ying Lu

**Author notes:** Corresponding authors: Ying Lu and Alfred L. Goldberg. These authors contributed equally to this work (Donghoon Lee, Yanan Zhu, Louis Colson).

## Abstract

Various hormones, kinases, and stressors (fasting, heat shock) stimulate 26S proteasome activity. To understand how its capacity to degrade ubiquitylated protein can increase, we studied ZFAND5, which promotes protein degradation during muscle atrophy. Cryo-electron microscopy showed that ZFAND5 induces large conformational changes in the 19S regulatory particle. ZFAND5’s AN1 Zn finger interacts with the Rpt5 ATPase and its C-terminus with Rpt1 ATPase and Rpn1, a ubiquitin-binding subunit. Surprisingly, these C-terminal interactions are sufficient to activate proteolysis. With ZFAND5 bound, entry into the proteasome’s protein translocation channel is wider, and ZFAND5 dissociation causes opening of the 20S gate for substrate entry. Using single-molecular microscopy, we showed that ZFAND5 binds ubiquitylated substrates, prolongs their association with proteasomes, and increases the likelihood that bound substrates undergo degradation, even though ZFAND5 dissociates before substrate deubiquitylation. These changes in proteasome conformation and reaction cycle can explain the accelerated degradation and suggest how other proteasome activators may stimulate proteolysis.

## Introduction

The 26S proteasome is the primary site of protein degradation in eukaryotic cells. It catalyzes the rapid hydrolysis of protein substrates marked for destruction by covalent attachment of one or more ubiquitin (Ub) chains. Protein ubiquitylation has been widely assumed to be the only regulated step in the ubiquitin-proteasome pathway and to be the only determinant of the substrate’s rate of degradation. However, it is now well established that the degradative capacity of the 26S proteasomes is also tightly regulated and increases in a variety of conditions when overall protein degradation in the cell increases, including starvation, heat shock, or muscle atrophy (Lee et al., 2018; Lee & Goldberg, 2022; VerPlank et al., 2019). Under these conditions, Ub conjugates accumulate, and thus proteasome function limits the rate of proteolysis. For example, during muscle atrophy, as may result from denervation, disuse, fasting or cancer, muscles express the Zn-finger protein, ZFAND5/ZNF216, which binds to 26S proteasomes and enhances their ability to hydrolyze ubiquitylated proteins, ATP, and small peptides ^1,2^. Proteasomes are also activated transiently (for 1-2 hours) upon phosphorylation by certain protein kinases, including DRK2 during the cell cycle ^3^ and Protein Kinase A or G in response to physiological challenges (e.g. exercise), hormones or neurotransmitters (e.g. epinephrine, glucagon, acetyl choline) that raise cGMP or cAMP levels ^4–6^, where proteasome function is impaired, drugs that raise cAMP or cGMP levels enhance proteasomal activity and can thus promote the clearance of the toxic aggregation-prone proteins causing these diseases ^4,6–8^.

The molecular mechanisms that account for the more efficient degradation of ubiquitylated proteins under these various conditions are completely unknown and have not been investigated previously. The degradation of a ubiquitylated substrate is a multistep, ATP-dependent process that involves a series of conformational changes, especially in the 19S regulatory particle (RP) ^9,10^. The ubiquitylated substrate may first bind directly to the proteasome through the Ub receptors (Rpn10, Rpn1, Rpn13) in the RP or bind indirectly via a “shuttling factor” ^11^. However, it is now clear that most ubiquitylated proteins that bind to proteasomes surprisingly dissociate without becoming committed to translocation into the 20S core particle (CP) for proteolysis ^12^. This poorly understood commitment step probably involves the capture of an unstructured region of the substrate by the hexameric AAA+ ATPase ring, which drives substrate translocation ^13^. These six ATPase subunits form a channel which during proteolysis become aligned with the gated pore in the CP’s outer α-ring ^11^. Opening of this gate is triggered by the ATPases’ conformational changes and is essential for substrate entry and degradation. Furthermore, to efficiently deliver a substrate into the CP, substrate-attached Ub chains must be removed by the proteasome-associated deubiquitylating enzymes (DUBs), Usp14, Uch37 and especially Rpn11. Despite appreciable progress in understanding these steps, it remains unclear which of these essential steps are rate-limiting and how their structural transitions are regulated to promote degradation. Increasing the proteasome’s degradative capacity likely requires an acceleration of one or more of these essential steps and a precise coordination of proteasome’s several structural transitions with the binding of the substrate molecule.

To understand how the 26S proteasome can be activated to degrade ubiquitylated substrates, we investigated the well-characterized proteasome activator ZFAND5, because of its importance in a variety of physiological processes requiring elevated capacity of protein breakdown. ZFAND5 expression is strongly enhanced in skeletal muscle during various types of atrophy (e.g. fasting, denervation, glucocorticoid treatment), where it is essential for the resulting rapid loss of muscle mass. Muscle atrophy occurs primarily through increased proteolysis by the ubiquitin-proteasome system (UPS) ^14^. ZFAND5’s capacity to stimulate hydrolysis of ubiquitylated proteins accounts for its ability to increase the overall rate of protein breakdown in cells ^1^ and explains why its induction is essential for this rapid loss of muscle mass ^2^.

In this study, we integrated cryo-EM structural determination with reaction kinetics analysis by single-molecule fluorescence microscopy, to define how ZFAND5 interacts with the 26S proteasome and activates it in substrate degradation. Our study shows that ZFAND5 reshapes the proteasome’s conformational landscape so as to increase the rate and efficiency of substrate degradation. The engagement of ZFAND5’s C-terminus with a regulatory site on Rpn1, induces a novel 19S conformation that appears to favor substrate translocation and proteolysis. Also the association of ZFAND5 through its A20 domain (amino acids 8-42) with the substrate and their simultaneous binding to the proteasome ensure that the 19S conformational changes are synchronized with the substrate arrival so as to maximize the stimulation of proteolysis. Surprisingly, ZFAND5’s rapid dissociation from the 26S also triggers key conformational changes that enable substrate translocation into the 20S CP. These findings not only expand our knowledge about the key steps in proteasome function, but also provide a conceptual and methodological basis for understanding the mechanisms of other proteasome activators.

## Results

### Defining the ZFAND5-proteasome interactions by cryo-EM and cross-linking mass spectrometry

To reveal the structural basis for the stimulation of proteasome activities by ZFAND5, we determined the cryo-EM structures of the human 26S proteasome complexed with recombinant ZFAND5. The presence of ZFAND5 significantly altered the conformational landscape of the proteasome. Unsupervised 3D classification identified eight distinct states—designated **Z+_A_** to **Z+_E_** and **Z‒_A_** to **Z‒_C_** with nominal resolutions of 3.6Å - 4.8Å (Fig. 1A; Fig. S1-S3). Five of them were designated Z+ states because they contain an extra density that can be fitted with an NMR structure of ZFAND5’s AN1 domain (amino acids 148-194) (PDB: 1WFL) which was extended using a structural prediction with AlphaFold ^15^. **Z+_A_** comprised 20.5% of the particles, **Z+_B_** 22.4%, **Z+_C_** 7.3%, **Z+_D_** 11.5%, and **Z+_E_** 7.2%. The other three states were named **Z‒**, because they lack any ZFAND5 density; **Z‒_A_** comprised 8.5% of the particles, **Z‒_B_** 8.5% and **Z‒_C_** 14.8%.

**Figure 1.**
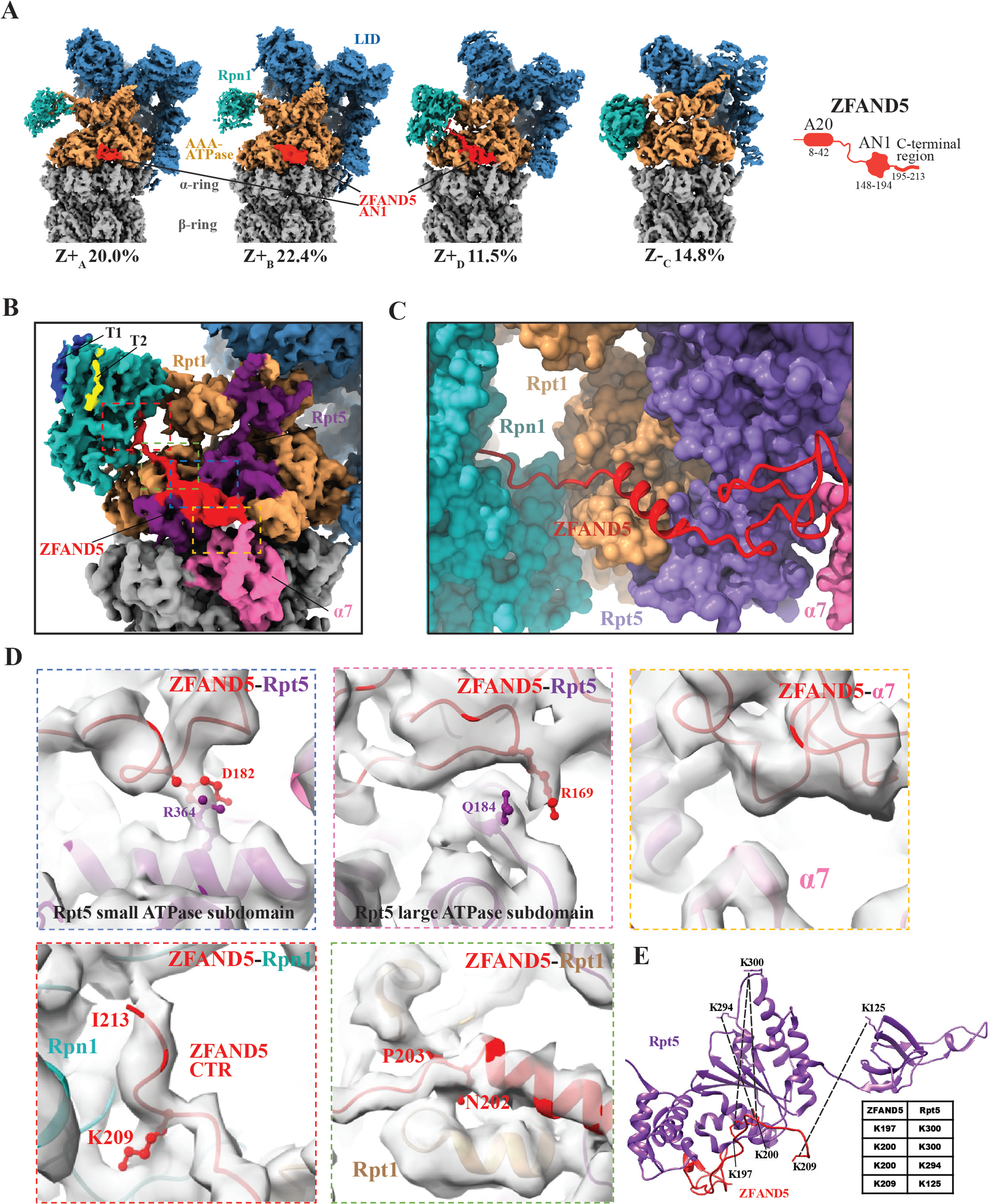
Cryo-EM structures of the human 26S proteasome complexed with ZFAND5/ZNF216. (A) Selected cryo-EM density maps of human 26S proteasomes in the presence of ZFAND5 (red). The relative abundances of each 3D class (% total). **Z+** refers to each class containing a ZFAND5 density and **Z‒** to classes lacking ZFAND5. As shown below, the domain architecture of ZFAND5 is shown on the right. (B) A local view of ZFAND5 in the cryo-EM density map of the **Z+_D_** state. Close-up views of ZFAND5’s interfaces with various proteasome subunits are shown in D, with regions of same color indicated in the top panel. (C) A local view of the **Z+_D_** model highlighting the molecular path that ZFAND5 follows on by Rpt5, Rpt1 and Rpn1 and the positions of its C-terminal residues in **Z+_D._** (D) Close-up views of ZFAND5’s interfaces with various proteasome subunits are shown in the lower panels with the same colors as in B. (E) Chemically-crosslinked residue pairs in a proteasome-ZFAND5 sample were identified by mass spectrometry and are represented as dotted lines.

In the **Z+** states, ZFAND5’s AN1 domain (residues R169 and D182) docks at the RP-CP boundary and interacts with both the large and the small domains of the ATPase subunit, Rpt5 (at residues Q184 and R364), and with the CP subunit, α7 (at residue V204) (Fig. 1B-D). To examine whether ZFAND5 may interact with additional sites that were not detected by cryo-EM, we cross-linked the ZFAND5-26S complex with disuccinimidyl sulfoxide (DSSO) which is reactive to primary amines and identified the cross-linked peptides, that were generated by trypsinization using mass spectrometry in a similar approach as used previously to define the detailed interactions between different proteasome subunits ^16^. Consistent with the cryo-EM result, lysine residues mainly in the C-terminus of ZFAND5 were cross-linked to lysines in the ATPase domain of Rpt5 (Fig. 1E and Table S1). No other 19S subunit was cross-linked to ZFAND5.

The **Z+_A_, _B_, _C_** states, which together comprised 54% of the particles, closely resemble the proteasome in the resting, or S_A_, state seen in the absence of ZFAND5 (RMSD=1.95Å) ^17^, with minor differences in the configurations of the non-ATPase subunits, Rpn1 and Rpn2 (Fig. S3E-S3I). The C-termini of Rpt3 and Rpt5 were found inserted into the inter-subunit pockets in the CP’s α-ring in these three states (Fig. S4A). These arrangements closely resemble those in the S_A_ state ^17^. As in that state the substrate translocation channel in the RP ATPases is not aligned with the gated entry channel in the CP. The **Z+_E_** state also resembles S_A_ or **Z+_A_**, except that its Rpn5 subunit only shows a partial density indicating structural flexibility (Fig. S4B). In these states, we did not resolve the regions that are N or C-terminal to ZFAND5’s AN1 domain, also likely due to structural flexibility of these regions.

### ZFAND5 C-terminal region interacts with Rpn1-Rpt1 in a novel RP conformation with an open substrate-translocation channel

The **Z+_D_** state represents a novel RP conformation that is distinct from either the resting or translocating states of the proteasome ^10,17,18^. Most subunits in the Lid subcomplex in **Z+_D_** exhibit a ∼5° clockwise rotation around the central axis, which causes these subunits to deviate by 5Å ∼10Å from their resting locations in the **Z+_A_** or S_A_ state (Fig. 2A). The arrangement of the ATPases, the pore loops and the status of nucleotide pockets in **Z+_D_** also differed from those in other states (Fig. S5 and S6).

**Figure 2.**
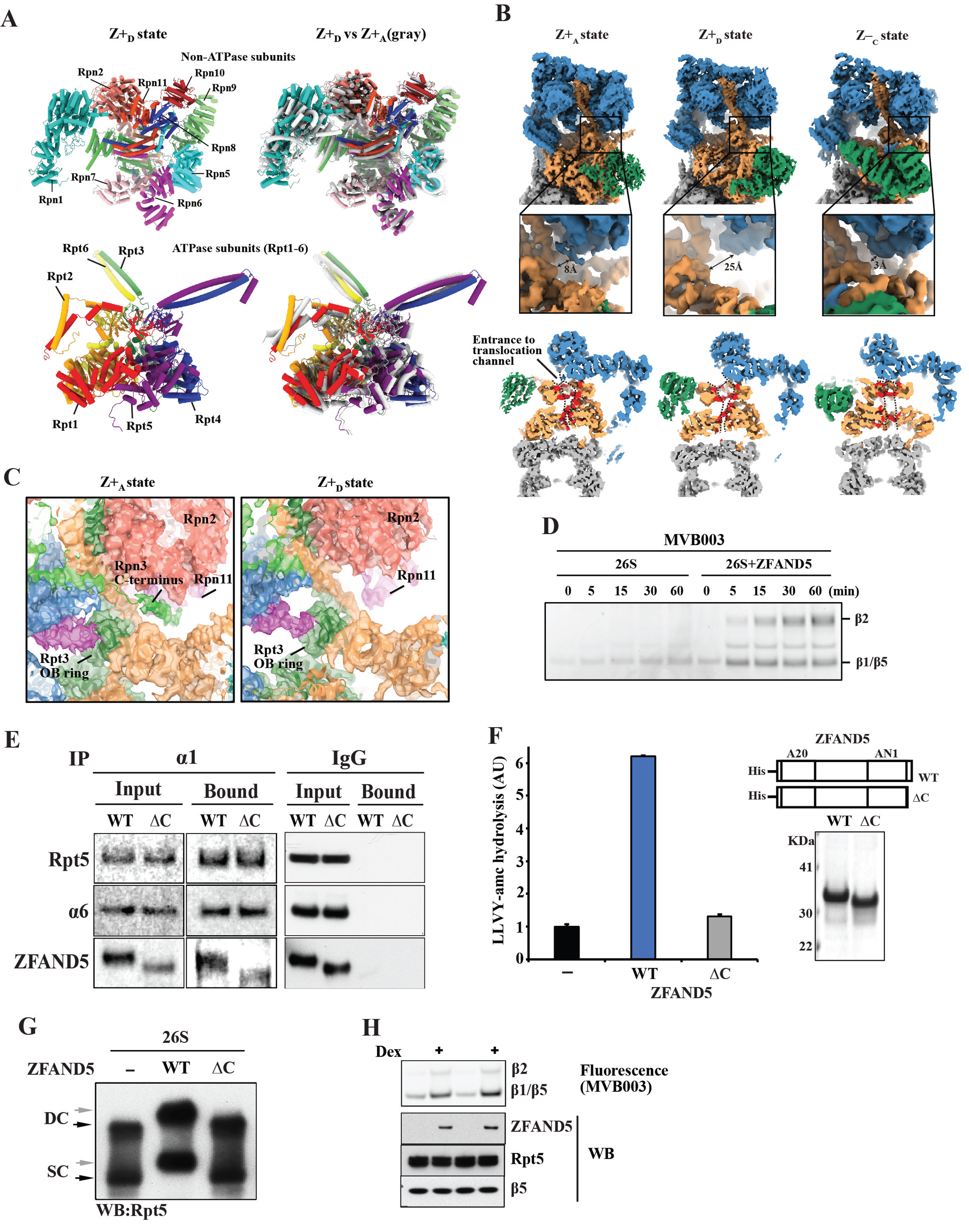
ZFAND5 induces a novel 19S conformation with an open entry to the substrate translocation channel in the Z+_D_ state. (A) Comparison of the structures of the ATPase and the non-ATPase parts in **Z+_D_** (left panel) and **Z+_A_** states. Superimposed on **Z+_D_** conformation (right panel) are shown in gray the locations of the corresponding subunits in **Z+_A_**, which are very similar in **Z+_B_** and **Z+_C_**. (B) Comparison of the entrance to the substrate translocation channel in three different 19S states. The diameter of the unobstructed part of the entrance is indicated by an arrow. Lower: vertical cross sections of cryo-EM maps. Substrate translocation channel’s interior residues are colored red, surface delineated by dashed lines. (C) Close-up views showing relocation of Rpn3’s C-terminal domain leading to unimpeded entry to the ATPase channel in **Z+_D_** states. (D) ZFAND5 increases access to active sites in the CP. An activity-based probe MVB003 for the active sites in the CP was incubated with 26S proteasome in the presence or absence of ZFAND5. Samples were analyzed by SDS electrophoresis and fluorescence imaging. (E) ZFAND5ΔC mutant lacking its 19 C-terminal residues binds to the 26S proteasome. Co-immunoprecipitation with immobilized anti-α1 antibody or control IgG. The bound samples were analyzed by Western blotting (WB). (F) The peptidase activity of 26S proteasome in the presence of ZFAND5 or its ΔC mutant. Right: Coomassie staining of purified ZFAND5. (G) Change in migration of proteasomes by ZFAND5 requires its C-terminus. 26S proteasomes were incubated with ZFAND5 or its ΔC mutant and were analyzed by native electrophoresis and by WB. (H) Induction of ZFAND5 by dexamethasone leads to proteasome gate opening shown by reactivity with MVB003. C2C12 myotubes were treated with dexamethasone (50mM) for 1 day and with MVB003 for 1h before harvesting. The labeled 20S subunits were resolved in SDS-PAGE, and proteasome contents were compared by WB.

The most striking feature of the RP in **Z+_D_** is its wide-open entrance to the substrate translocation channel, a rigid ring formed by the OB domains of the ATPases, which is partially occluded in the resting states, such as **Z+_A_** and S_A_ ^17,19^. The diameter of the unobstructed portion of the entrance to the channel in **Z+_D_** is 25Å compared to 8Å in the other **Z+** states presumably as a result of the Lid rotation (Fig. 2B). Opening of the translocation channel probably also results from the displacement of a helix domain of Rpn3 (Fig. 2C). This helix is sandwiched and is stabilized by Rpn11 and the OB domain of Rpt3, and rests above the OB ring, so as to occlude the translocation entrance in the S_A_ state (Fig. 2C).

In the **Z+_D_** map, a density that corresponds to the C-terminal region (CTR, amino acids 195-213) of ZFAND5 is clearly visible, which is not evident in the **Z+_A_, _B_, _C_** states. This density extends from ZFAND5’s AN1 domain, loops over and arrives at a cavity that is adjacent to Pro719 in the convex side of the toroidal domain of the non-ATPase subunit Rpn1 and is also flanked by the OB domain of the ATPase Rpt1 (Fig. 1C). This binding site on Rpn1, which we term the Z site, was previously not known to be involved in protein-protein interactions and is separate from the T1 and T2 sites on Rpn1, which are located on the opposite side of Rpn1 and bind Ub and UBL-domain-containing proteins (Fig. 1D) ^20^. The functional importance of these interactions is studied below.

In **Z+_D_**, ZFAND5’s CTR forms additional contacts with the RP and is likely stabilized by these interactions. A short helix emerges adjacent to ZFAND5’s AN1 (residues N202 and P203) and interacts with another ATPase Rpt1 through helix 5 in Rpt1’s AAA domain (at residues V202 and N314) (Fig. 1C and 1D). Beyond Rpt1, ZFAND5’s residues are also engaged in specific interactions with a C-terminal fold of Rpn1, which may help position ZFAND5’s C-terminus (the residues K209 and I213) to the Z site in the toroidal cavity of Rpn1 (at residues P719 and V850) (Fig. 1D and 1E). To accommodate ZFAND5, a Rpn1 helix (Asp346-Gly360) that interacts with Rpt1 in the S_A_ state is displaced and joins the toroid of Rpn1 in **Z+_D_**. This change may destabilize Rpn1-Rpt1 interaction and causes a ∼30° rotation of Rpn1 away from Rpt1 in **Z+_D_** (Fig. S7A). Interestingly, the local resolution of Rpn1 in **Z+_D_** is significantly higher than that in the other states, indicating that docking of ZFAND5’s CTR probably stabilizes the configuration of Rpn1 in the RP (Fig. S7B). These residues involved in ZFAND5’s CTR-19S interactions are highly conserved in high eukaryotes (Fig. S8).

### Upon dissociation ZFAND5 promotes translocation-competent proteasomal states

Translocation of the protein substrate into the CP requires both the opening of the gated pore in its α-ring and the alignment of the 19S ATPase channel with this open gate. Gate opening in the CP is controlled by conformational changes of the ATPases ^9,10,21,22^. Surprisingly, the CP gate is in a closed conformation in all the **Z+** states that contain a ZFAND5 density, but it adopts an open conformation in all the **Z‒** states (Fig. S4A). In addition, in all these **Z‒** states, the substrate translocation channels in the RP and the CP are aligned, which is a characteristic feature of proteasomes active in degradation ^9,10,19^. The **Z‒_A_** and **Z‒_C_** states resemble respectively the translocation-competent E_D1_ and E_D2_ structures of the substrate-engaged human proteasome (or 5D and 4D for yeast 26S), with respect to both the RP geometry and the nucleotide-binding status, although no ubiquitylated substrate was included in the cryo-EM sample (Fig. S6, S7C and Table S2)^10^. ZFAND5 was not degraded by the 26S, and its level was stable during the incubation with the proteasome (Fig. S9A and S9B).

The exposure to ZFAND5 dramatically enlarged the population of open-gated particles, which increased from 7.9% in the 26S sample without ZFAND5 ^17^ to 31.8% in its presence (Table S2). Substrate translocation into the CP requires an open gate (i.e. a **Z‒**) conformation. Because ZFAND5 was not present in these particles, the transition of the proteasomes into a translocation-competent, **Z‒** state must occur upon ZFAND5 dissociation. Direct evidence for this conclusion is shown below.

Further evidence for the increase in gate-opening was obtained using an activity-based probe (yellow Bodipy-Cy3-epoxomicin, MVB003). This agent covalently modifies the active sites within the 20S core particle ^23^, and thus rapid derivatization of these subunits is a measure of their accessibility. Exposure of proteasomes to ZFAND5 dramatically increased their covalent modification (Fig. 2D and S9C), as expected from the large increase in open-gated particles.

This increase in the open-gated population can account for the large stimulation of peptide hydrolysis by ZFAND5 seen previously ^1^. This stimulation requires the AN1, but not the A20 domain of ZFAND5. To test the dependence on ZFAND5’s CTR, we truncated ZFAND5’s C-terminus by 19 amino acids (ZFAND5ΔC). ZFAND5ΔC maintained the interaction with the proteasome in a co-precipitation assay (Fig. 2E), but it could not stimulate peptide hydrolysis (Fig. 2F). In addition, both the doubly- and singly-capped proteasomes migrated more slowly in native-gel electrophoresis after exposure to ZFAND5, which probably reflects the large conformational changes in the RP ^1^. However, neither ZFAND5ΔC nor ZFAND5 with a mutated AN1 domain slowed proteasome migration in native PAGE (Fig. 2G and S9D). Therefore, the activation of the 26S proteasome through transition into the open-gated **Z‒** states requires ZFAND5’s CTR. In addition, because the CTR does not appear to interact with proteasome in the **Z+_A,B,C,E_** states, transition into **Z‒** presumably follows the formation of the **Z+_D_** state.

We then tested whether ZFAND5 expression caused a similar increase in the open-gated population of intracellular proteasomes, as was shown with purified particles (Fig. 2D). C2C12 myotubes were incubated for one day with dexamethasone, which induces ZFAND5, and then were exposed for 1h with the membrane-permeant fluorescent probe MVB003. Although the myotube content of 19S (Rpt5) or 20S (β5) particles did not change with dexamethasone, the reactivity of 20S active sites increased dramatically (Fig. 2H). Thus, in cells, ZFAND5 seems to increase the fraction of particles in an open-gated conformation, as observed with purified particles.

### ZFAND5 accelerates proteasomal degradation of ubiquitylated substrates

In order to understand the functional effects of ZFAND5-induced structural changes, we next studied the degradation of ubiquitylated proteins in the presence of ZFAND5. We used a fluorescent model substrate containing the N-terminal region of cyclinB (cycB) fused to a destabilized fluorescent protein cpGFP ^24,25^, and studied how proteasome activation depends on ZFAND5’s structural domains. Ubiquitylation of cycB-cpGFP by the E3 Anaphase-Promoting Complex (APC/C) leads to efficient and processive degradation by purified 26S proteasomes (Fig. 3A), with little deubiquitylation or partial cleavage of the substrate (Fig. S10A). This reaction appeared to obey the Michaelis-Menten kinetics, and a Lineweaver–Burk analysis indicated an apparent K_M_ of 5.6nM for this substrate (Fig. S10B). The turnover time for this substrate was 46 seconds/substrate/proteasome at substrate concentrations above the K_M_. Most of our subsequent experiments were performed under multiple-turnover conditions with a large substrate excess.

**Figure 3.**
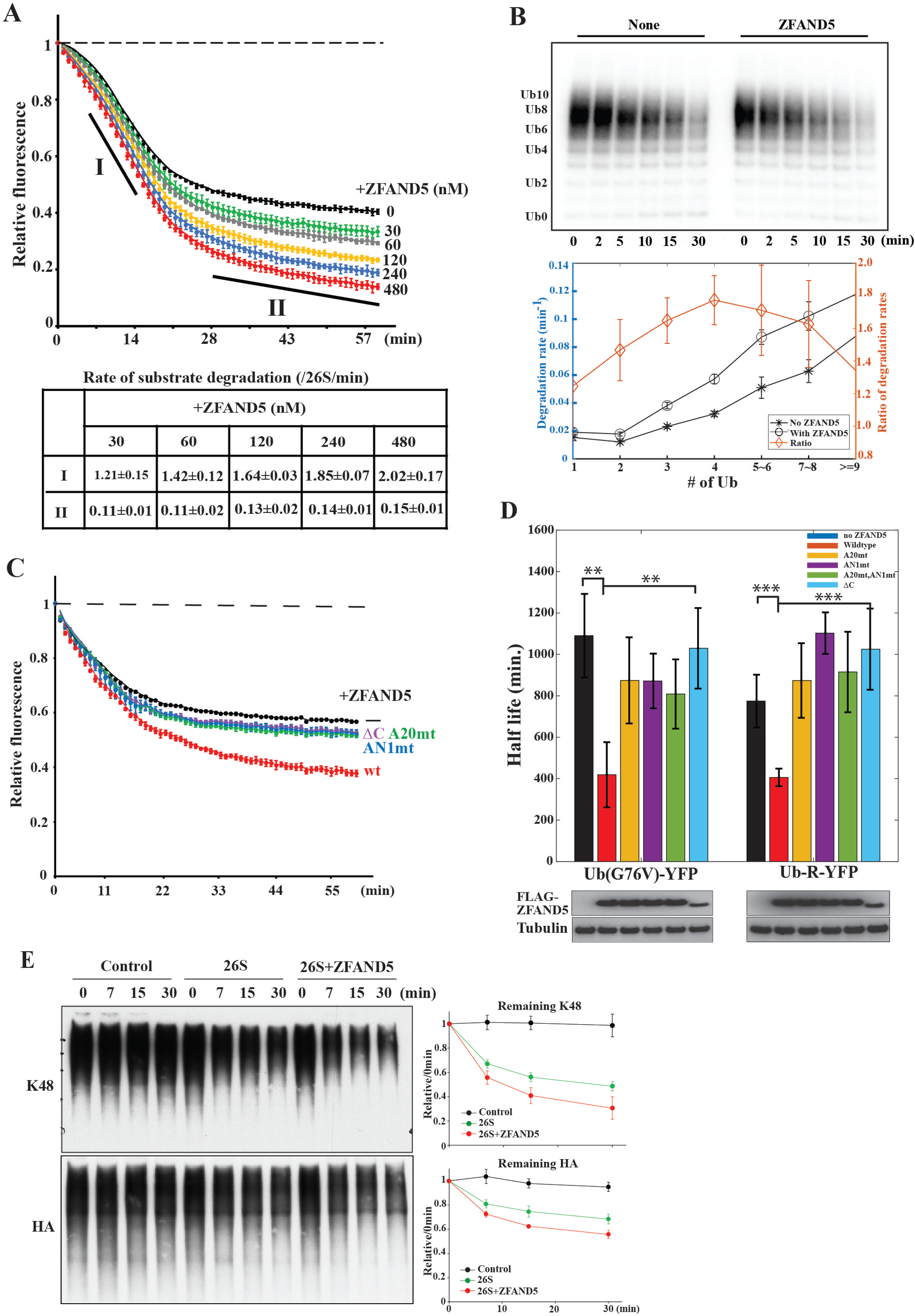
ZFAND5 stimulates the degradation of Ub conjugates by the 26S proteasome. (A) Intensities of a fluorescent substrate, ubiquitylated cycB-cpGFP, upon degradation reactions with purified 26S proteasomes and ZFAND5 at various concentrations. CycB-cpGFP’s degradation rates calculated from the slopes at the initial (“I”) or late (“II”) stage of the reaction are listed in the table. Shown are the mean rates ± the standard deviations (SD) of five replicates. The degradation rates for the indicated time frames (I and II) are shown ± SD from three independent experiments. (B) Effect of ZFAND5 on the degradation of substrates containing increasing numbers of Ub molecules. ^32^P-labeled ubiquitylated cycB-cpGFP was incubated with 26S proteasomes and ZFAND5. Samples were analyzed by autoradiography and the degradation rate of each ubiquitylated species was plotted on the right. Error bars represent uncertainty in quantification. (C) ZFAND5’s C-terminal 19 residues, A20 domain and AN1 domain are essential for the stimulation of degradation. Reactions were performed as in A but with ZFAND5 mutants. Error bars represent the SD of five replicates. (D) Effects of ZFAND5 and its mutants on the mean half-lives of, Ub-R-YFP, a substrate of the N-end rule pathway, and Ub- (G76V)-YFP, a substrate of the UFD pathway in cells. Degradation of these substrates was monitored using time-lapse microscopy in a cycloheximide chase experiment in HEK293 cells cotransfected with ZFAND5 WT or mutants. Cells expressing 5∼15uM reporters were selected. 15∼50 cells were analyzed in each category. Error bars represent the standard errors and the statistical significance is marked by “*”. The expression levels of FLAG-ZFAND5WT and mutants were examined by WB. (E) ZFAND5 stimulates the degradation of naturally ubiquitylated proteins, which were isolated from HEK293 cells expressing polyHis-HA-tagged Ub, followed by incubation with purified 26S proteasome with or without ZFAND5. The samples were analyzed for K48 Ub linkage and HA epitope by WB. Error bars represent the SD of three replicates.

Degradation of ubiquitylated cycB-cpGFP proceeded in different phases. The apparent degradation rate was slower in the late phase (“II”, 5.9 minute/substrate) than in the initial phase (“I”, 0.59 minute/substrate) (Fig. 3A). ZFAND5 increases the initial degradation rate by 24%, but the late phase of the reaction by 71% (Fig. 3A) with Ka ∼100nM (Fig. S10C). Preincubation of ZFAND5 with either the substrate or the proteasome further increased its stimulatory effect (Fig. S10D). This lower degradation rate in phase II is unlikely to be due to substrate depletion, because the remaining substrate concentration measured by GFP intensity was still much higher than the K_M_ of 5.6nM (Fig. S10B), nor was it caused by a loss of proteasome activity during incubation, since we recorded an almost identical degradation rate by adding a second dose of ubiquitylated cycB-cpGFP during phase II of the reaction (Fig. S10E). Also, ZFAND5 is not consumed during these incubations with the proteasome and the ubiquitylated substrate (Fig. S10F).

One contributor to this multi-stage kinetics is substrate heterogeneity. The Ub conjugates formed on cyclin-B by APC/C vary in their configurations of Ub moieties ^26^ which determine their susceptibility to proteasomal degradation ^12^. Since the Ub copy number is a key determinant of the degradation rate ^12^, we measured the rates of degradation of cycB-cpGFP conjugated with different numbers of Ub moieties. While ZFAND5 increased the hydrolysis of all ubiquitylated cyclinB species, substrates containing 4-6 Ub moieties were stimulated to the greatest extent (Fig. 3B). Thus, ZFAND5 appears to stimulate the degradation of substrates carrying a range of Ub stoichiometry normally preferred by proteasomes ^27^ but it seems to have a larger effect on substrates with intermediate Ub copies that are suboptimal for proteasome recognition.

Inactivating either the Zn finger domains of ZFAND5, the AN1 domain or the Ub-binding A20 domain, largely abolished the stimulation of the degradation of cycB-cpGFP (Fig. 3C) ^1^. Also, as found for peptide hydrolysis (Fig. 2F), ZFAND5’s CTR is required for the enhanced cycB degradation (Fig. 3C). While the A20 domain must function in Ub conjugate binding, the requirement for the C-terminus and AN1 domain implies that their interactions with the Rpn1, Rpt1 and Rpt5 in the **Z+_A_** to **Z+_D_** conformations are also critical for the accelerated proteolysis.

### ZFAND5 accelerates the degradation of model UPS substrates in cells

In MEF cells lacking ZFAND5 the overall rate of degradation of cellular proteins is reduced, and its induction appears essential for the rapid acceleration of overall proteolysis during muscle atrophy ^1,2^. However, ZFAND5’s effects on the degradation of specific proteins in cells have not been investigated. Therefore, we examined the effects of ZFAND5 expression on the half-lives of two well-studied model substrates of the UPS, Ub-R-YFP, a substrate of the N-end rule pathway, and of Ub(G76V)-YFP, which is ubiquitylated by the Ub-fusion-degradation (UFD) pathway. These constructs were co-expressed with WT or ZFAND5 mutants in HEK293T cells, and we used time-lapse microscopy to quantitatively study their degradation in individual cells after addition of cycloheximide to block protein synthesis.

In cells expressing these reporters at high levels (5∼15uM), ZFAND5 markedly shortened the average half-life of both (Fig. 3D). Importantly, their degradation was not accelerated upon expression of ZFAND5 mutants lacking the A20 or AN1 domains or its C-terminus, as was found for the degradation of cycB fusions proteins by purified proteasomes (Fig. 3D). Because ZFAND5 is not known to interact with other factors, such as the E3s, that are required for the degradation of these reporters, the most likely explanation for this result is that ZFAND5 in cells stimulates the degradation of these reporters by activating the proteasome, through the same mechanisms studied *in vitro.* However, co-expressing ZFAND5 did not significantly alter the stability of these proteins in low expressing cells, presumably because the rate of reporter degradation is not limited by the proteasome activity.

To learn if ZFAND5 can also stimulate the degradation of the diverse Ub conjugates found *in vivo*, we isolated Ub conjugates from growing HEK293 cells stably expressing polyHistidine-HA tagged Ub and added N-ethylmaleimide to inhibit hydrolysis of Ub conjugates by DUBs. When these proteins ubiquitylated in the cell were incubated with purified proteasomes in the presence of ZFAND5 for 30min, we observed a stimulation of Ub conjugate degradation, as indicated by the greater decrease in K48 Ub chains (Fig. 3E). This capacity to increase degradation of ubiquitylated proteins presumably accounts for the enhancement by ZFAND5 of degradation of endogenous proteins in cells and in crude extracts ^1^.

### ZFAND5 promotes the degradation of difficult-to-unfold substrates

The rate of substrate degradation by the proteasome is limited not only by the nature of the Ub modification but also by structurally stable domains that exist in many cellular proteins. Substrates with these types of domains, after proteasome binding, may resist unfolding, deubiquitylation or proteolysis and may cause incomplete (non-processive) degradation with the release of partially digested fragments ^28^. To study the effect of ZFAND5 on the degradation of such hard-to-unfold substrates, we fused cycB with the fluorescent protein EGFP, which the 26S proteasome by itself degrades much more slowly than cpGFP (as measured by the loss of the fluorescence signal), even when their levels of ubiquitylation were similar (Fig. S11A) ^25^. However, in the presence of ZFAND5 the degradation rate of the EGFP moiety was stimulated by 70% which was greater than the fold change seen with cycB-cpGFP (although the absolute degradation rate of the hard-to-unfold substrate was still slower than that of cpGFP) (Fig. 4A and S11B).

**Figure 4.**
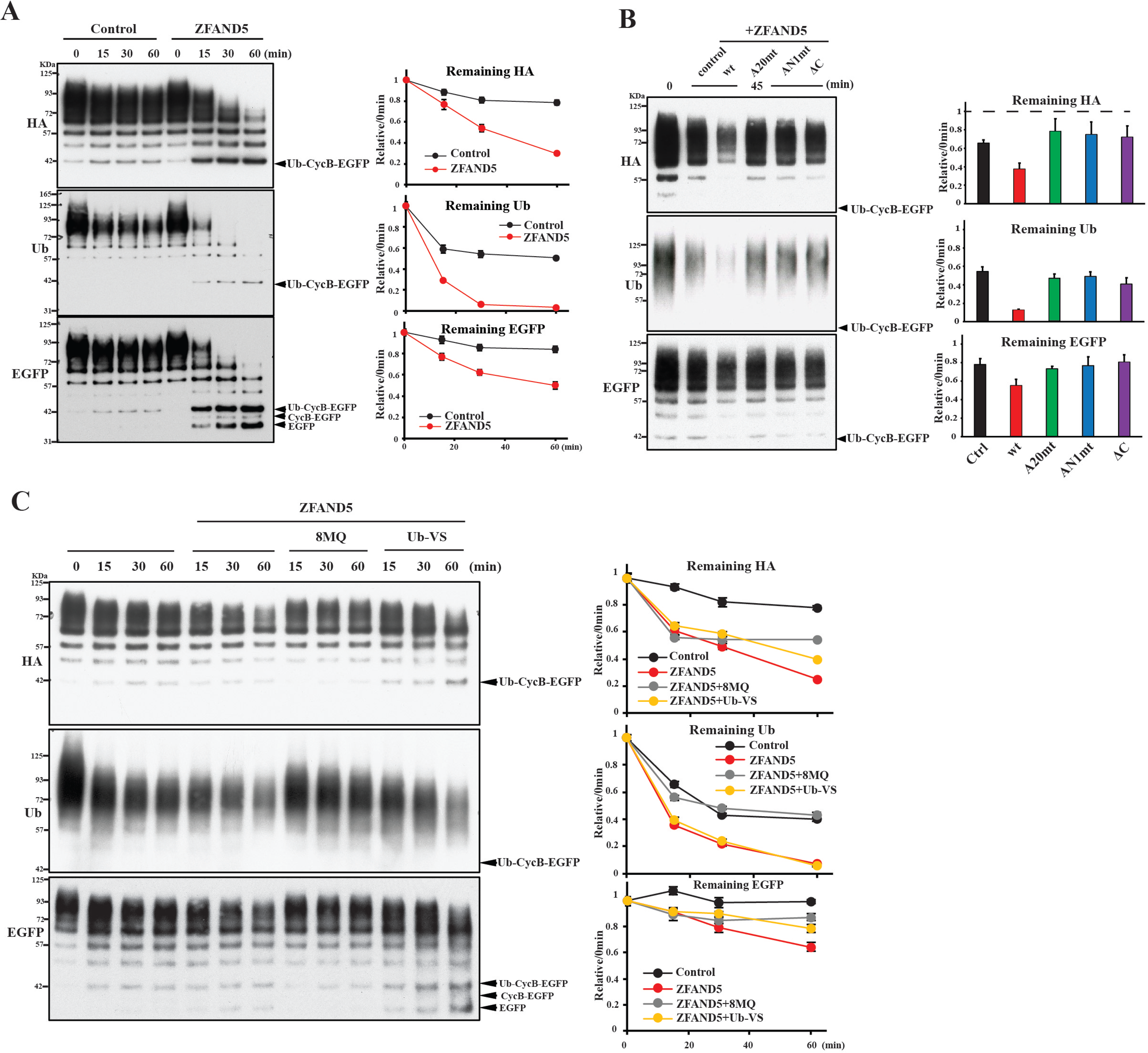
ZFAND5 stimulates deubiquitylation and degradation of unfolding-resistant substrate by the 26S proteasome. (A) Ubiquitylated cycB-EGFP was incubated with 26S proteasome in the presence of WT and mutant ZFAND5 in (B). Levels of cycB, EGFP and ubiquitylation were determined by WB. Error bars represent the SD of three replicates. (C) Stimulation of degradation and deubiquitylation requires Rpn11. Degradation assays were performed as in (A), but in the presence of the cysteine DUB inhibitor Ub-VS or the Rpn11 inhibitor 8MQ. Error bars represent uncertainty in quantification.

A western blot analysis of the products released by the proteasome indicated that ZFAND5 greatly accelerates both the deubiquitylation and non-processive degradation of cycB-EGFP as well as its complete hydrolysis, which was dependent on ubiquitylation of the substrate (Fig. 4A and S11C). The stimulation of all these proteasomal activities also requires ZFAND5’s AN1, A20, and C-terminal domains (Fig. 4B). In the presence of ZFAND5, removal of the Ub chains from this substrate was complete within about 30 minutes, which is much faster than the deubiquitylation rate without ZFAND5. As expected ^29,30^, deubiquitylation of cycB-EGFP was blocked by the Zinc chelator, 8-mercaptoquinoline (8MQ), but was not affected by Ubiquitin Vinyl Sulfone (Ub-VS), an inhibitor of cysteine DUBs (Fig. 4C). Thus, this process is driven mainly by the metalloprotease Rpn11, which resides at the entrance to the ATPase channel. Inhibition of Rpn11 did not prevent ZFAND5’s stimulation of the degradation of the peptide substrate (Fig. S11D).

Processing of difficult-to-unfold substrates can lead to proteasome inhibition through the formation of stable (non-degraded) intermediates ^29,31^. In these experiments, both deubiquitylation and partial degradation significantly slowed down after 10 minutes in the absence of ZFAND5, probably because of proteasome inactivation by some substrate molecules (Fig. 4A). In contrast, degradation of the hard-to-unfold substrate did not terminate rapidly when ZFAND5 was present, which suggests that the ZFAND5-activated proteasomes resisted such inhibition.

### A peptide corresponding to ZFAND5’s C-terminal residues by itself can stimulate proteasome activities

Both our structural and mutation studies suggest the functional importance of ZFAND5’s C-terminal residues. To further explore the activity of ZFAND5’s CTR, we synthesized a peptide corresponding to residues 195-213 and tested whether by itself this 19-residue peptide might influence proteasome activities. Surprisingly, the addition of this peptide alone stimulated peptide hydrolysis 2.5-fold (Fig. 5A) and also enhanced the hydrolysis of ubiquitylated cycB, primarily in the initial phase of the degradation reaction (Fig. 5B). These stimulatory effects were smaller than those of full-length ZFAND5, which increased peptide hydrolysis by up to 10-fold and stimulated cycB degradation in both phase I and II (Fig. 3A). By contrast, if a 19-residue peptide from an unrelated protein was added, proteasome activity did not increase. Interestingly, we found that N-terminal acetylation of the CTR peptide also attenuated the stimulation. By contrast, neither the isolated A20 nor the isolated AN1 domain of ZFAND5 could enhance the proteasome’s peptidase activity ^1^. Then, we examined if CTR peptide binds to the same site and competes with 26S-bound ZFAND5. When the CTR peptide was present at the same concentration as that enhances the 26S peptidase activity, the proteasome-bound full-length ZFAND5 clearly reduced, suggesting that the CTR peptide and ZFAND5 compete for binding to the 26S (Fig. S12). These findings make it very likely that the **Z+_D_** state, in which ZFAND5’s CTR interacts with Rpn1 and Rpt1, is a key intermediate in proteasome activation, and that these interactions are both necessary and sufficient to induce many of the conformational changes leading to gate opening and enhanced proteolytic activity.

**Figure 5.**
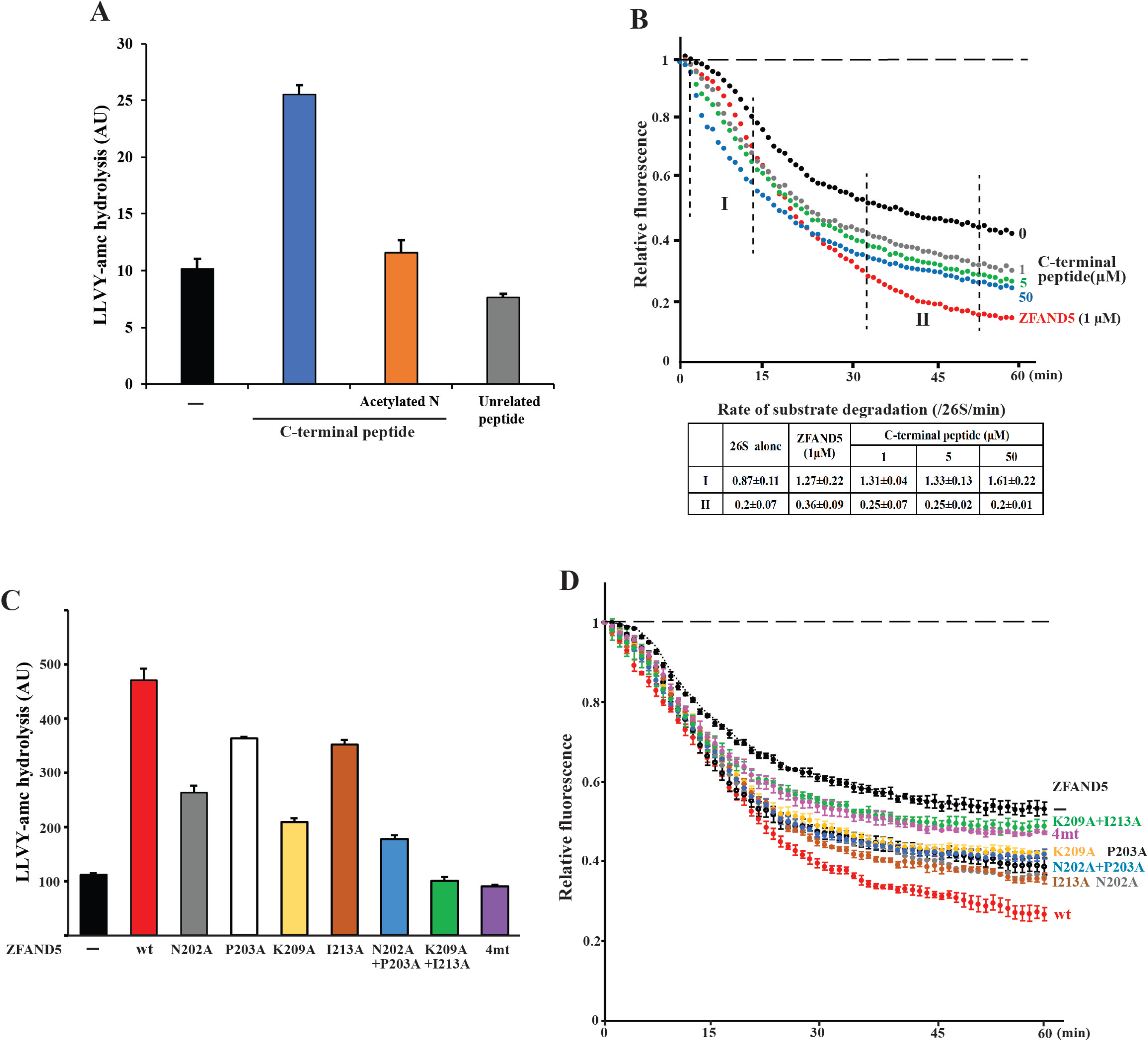
A 19-residue peptide from ZFAND5’s C-terminus by itself activates proteasomal degradation of short peptides and Ub conjugates. (A) The chymotryptic peptidase activity of purified 26S proteasome was determined in the presence of indicated peptides. Error bars represent the SD of three replicates. (B) The CTR peptide stimulates degradation of ubiquitylated substrates. Increasing concentrations of the CTR peptide were incubated with ubiquitylated cycB-cpGFP and 26S, and degradation was measured by fluorescence intensity. The degradation rates during the indicated time frame are presented ± SD from three independent experiments. (C) Mutagenesis of the proteasome-interacting residues of ZFAND5 and the effects on proteasome activation. The peptidase activity was determined as in A, but with indicated ZFAND5 variants. (D) As in B, but in the presence of indicated ZFAND5 variants. Results are presented as the mean ± SD of three replicates.

### The interactions of ZFAND5’s C-terminal residues with the RP are important for proteasome activation

To further investigate the role of ZFAND5’ CTR interactions with Rpn1 and Rpt1 in proteasome activities, we examined the effects of the specific residues’ interactions by site-directed mutation. A total of seven mutations, single, double or four residues to alanine, where four residues apparently interact with Rpn1 (i.e. K209 and I213) or Rpt1 (i.e. N202 and P203) were made. Intriguingly, mutation of individual residues caused a decrease in the 26S peptidase activity and hydrolysis of the ubiquitylated substrate (Fig. 5C and 5D). Double mutations at the residues interacting with Rpn1 or Rpt1 further lowered the stimulatory effect of ZFAND5, and mutations at all four residues blunted the stimulation of both peptidase and the degradation of Ub conjugates. It is noteworthy that the effect of loss of the Rpn1 interactions (i.e. both K209A and I213A) appears as strong as the mutations at all four residues. This result confirms that the conformational changes induced by the CTR-RP interactions are critical for the stimulation of 26S activity.

### Single-molecule analysis of ZFAND5-substrate-proteasome interactions

To further understand the mechanism of proteasome activation by ZFAND5, we investigated the kinetics of the key steps in the degradation process using single-molecule fluorescence microscopy. We first examined the ZFAND5-proteasome interactions. Purified 26S proteasomes were immobilized on a passivated glass surface via an anti-20S antibody (Fig. 6A). The binding of ZFAND5, which had been fluorescently labeled using an N-terminal SNAP tag, was monitored by Total Internal Reflection Fluorescence (TIRF) microscopy. The presence of the SNAP-tag did not affect the ability of ZFAND5 to stimulate the proteasome’s peptidase activity (Fig. S13A).

**Figure 6.**
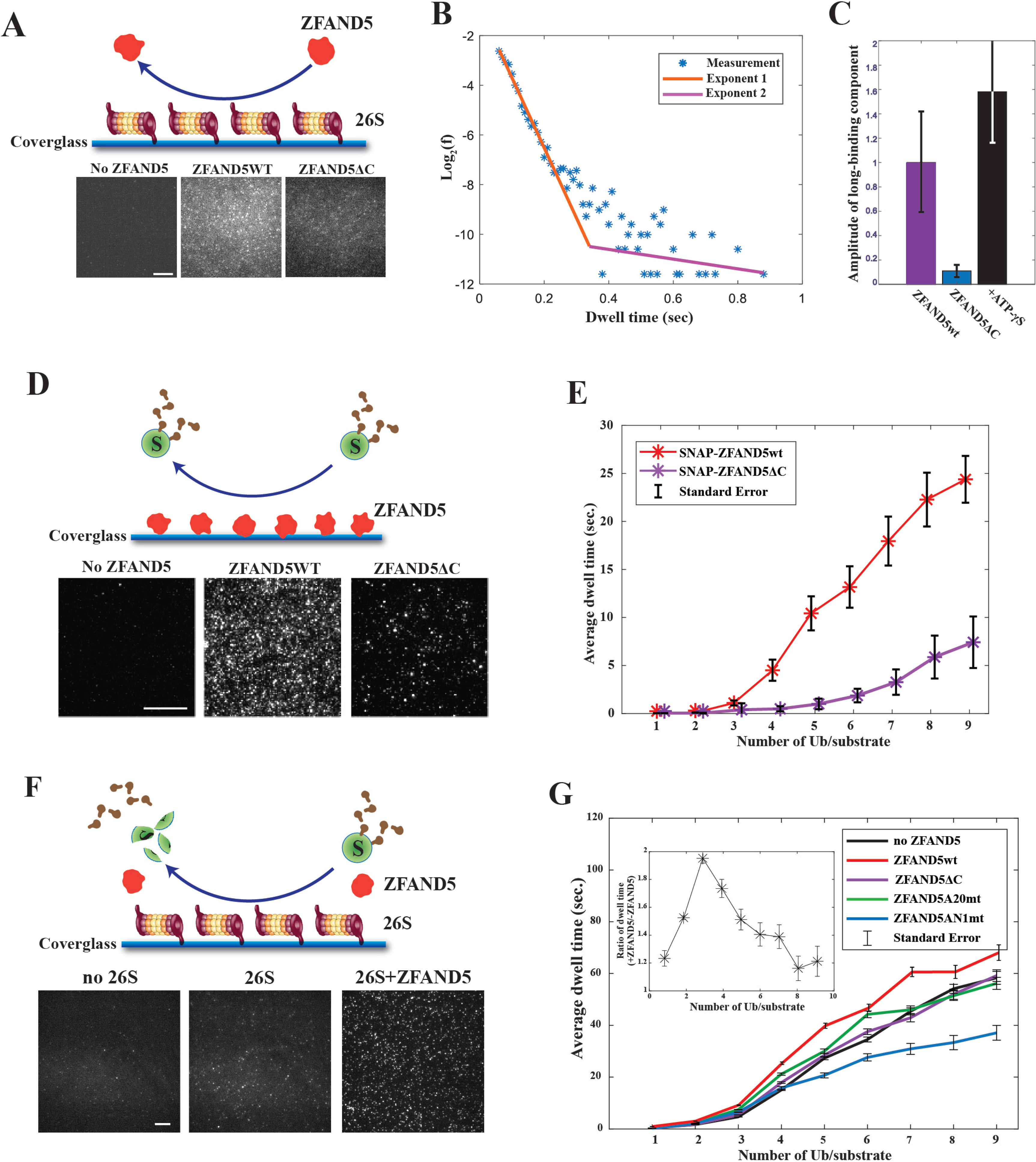
ZFAND5 exhibits two binding modes on the 26S proteasome and enhances substrate-proteasome interaction. (A) ZFAND5’s C-terminal region is essential for the stimulation of substrate association with the proteasome. Schematic and typical images of the single-molecule assay testing the interaction of fluorescent ZFAND5 with surface-immobilized 26S particles. Scale bars = 5µm (B) Distribution of ZFAND5’s dwell times on the proteasome. The measurements were fitted with a double exponential function and the two exponents are plotted as straight lines on a semi-log scale. (C) Amplitude of the second (i.e. long-binding mode) exponential component of ZFAND5’s dwell time distribution. Measurement and data processing were performed as in B in the presence of ZFAND5, or ZFAND5ΔC, or upon addition of ATP-γS. Error bars represent fitting uncertainty. (D) Schematic and sample images of the single-molecule assay to study the interactions of securin conjugated with Dy550-Ub and immobilized ZFAND5 WT or ΔC. The average dwell-time of a securin molecule on ZFAND5 is shown in (E). (F) Schematic and sample images of the single-molecule fluorescence analysis of the kinetics of substrate processing by immobilized 26S proteasomes. (G) The average dwell-time of a securin molecule conjugated with Dy550-Ub on the 26S proteasome in the presence of 500µM ZFAND5 or its mutants was plotted vs. the number of Ub molecules per substrate molecule. Inset: the ratio of the dwell-time values in the presence and absence of ZFAND5.

ZFAND5 associated transiently with the proteasome. The distribution of ZFAND5’s dwell time (∼1/k_off_) on the proteasomes indicated at least two exponential modes, with time constants of 50±15ms and 614±350ms respectively (Fig. 6B; Fig. S13B). Based on our Cryo-EM analysis, we hypothesized that engagement of ZFAND5’s CTR with Rpt1 and Rpn1 may stabilize the ZFAND5-proteasome interaction and account for the long-binding mode. Accordingly, deletion of the 19 C-terminal residues of ZFAND5 reduced the long-binding component by about 10X (Fig. 6C; Fig. S13C). Thus, the ZFAND5-proteasome interaction is stabilized by the CTR, most likely through its engagement with Rpn1 and Rpt1. Interestingly, the addition of ATP-γS promoted the long-binding mode, suggesting that efficient dissociation of ZFAND5 from the proteasome requires ATP hydrolysis, a process that is also stimulated by ZFAND5 ^1^.

We next examined ZFAND5’s affinity for ubiquitylated substrates. ZFAND5 was directly immobilized onto the surface via the SNAP tag and was incubated with polyubiquitylated securin (Fig. 6D), in which the Ub molecules were fluorescently labeled, so that the number of Ubs on the substrate molecule could be determined by the fluorescence intensity of the substrate spot ^12^. The substrate’s dwell time on the surface increased with the increase of the Ub stoichiometry, suggesting an avidity effect in the interaction of ZFAND5 with the ubiquitylated protein (Fig. 6E).

The A20 domain on ZFAND5 and other proteins is essential for binding Ub chains ^32^. Surprisingly, ZFAND5’s C-terminal residues also appear to be directly involved in the substrate interaction, since the dwell time of ubiquitylated securin on the surface-immobilized SNAP-tagged ZFAND5ΔC was much shorter than that of WT ZFAND5 (Fig. 6E). Because this CTR lacks a recognizable Ub-interacting motif, the molecular basis for this effect on substrate binding is still unclear.

The addition of ZFAND5 increased the total amount of substrate-bound proteasome particles. This effect was at least in part due to prolongation of the substrate’s dwell time on the proteasome (Fig. 6F and 6G). Ub conjugates showed a similar dwell time on the surface-immobilized proteasomes as on ZFAND5. ZFAND5 appeared to increase most the dwell time of substrates bearing 3-5 Ubs, which correlates with the optimal Ub stoichiometry for the stimulation of proteolysis by ZFAND5 (Fig. 3B). As expected, the Ub-interacting A20 domain of ZFAND5 is essential for the dwell time enhancement, as a mutation inactivating the A20 Zn finger largely abolished this effect, without affecting the ZFAND5-proteasome interaction (Fig. 6G). Interestingly, mutating the AN1 domain resulted in an even shorter dwell time than that seen with the Ub conjugates in the absence of ZFAND5. Presumably this decrease in dwell time contributes to the inability of these mutants to stimulate Ub conjugate degradation (Fig. 3C). The CTR of ZFAND5, which is essential for stimulating Ub conjugate degradation, is also required for the dwell-time enhancement. Since the proteasome interacts with ZFAND5 more transiently than with the substrate, ZFAND5 prolongs the substrate’s dwell time most likely through altering the proteasome’s conformations, rather than by of continuing to function in a substrate-ZFAND5-26S complex (e.g. as an alternative 19S substrate receptor).

### ZFAND5 increases the likelihood of a proteasome-bound substrate undergoing degradation

Most of the ubiquitylated substrates that bind to the 26S proteasome dissociate without degradation ^12^. We therefore examined if ZFAND5 would affect the likelihood that a proteasome-bound substrate will undergo processive deubiquitylation and proteolysis. To this end, we used the rapid and ‘stepped’ decrease of Ub intensity in single-molecule traces as a signature for substrate deubiquitylation and translocation (Fig. 7A)^12^. The cycB substrate contains multiple Ub chains formed by the APC/C. These chains are deconjugated co-translocationally by Rpn11 (once another 26S DUB Usp14 has been depleted from proteasome), resulting in ‘stepped’ Ub signal reductions which typically are completed within 10∼20 seconds. In the absence of ZFAND5, only a small fraction (∼14%) of substrate-proteasome encounters resulted in deubiquitylation and translocation, as was observed previously (Fig. 7B) ^12^. In contrast, ZFAND5 increased the fraction of such events from 14% to 23%, and this increase in the likelihood of degradation required the presence of ZFAND5’s CTR and its A20 and AN1 domains (Fig. 7B).

**Figure 7.**
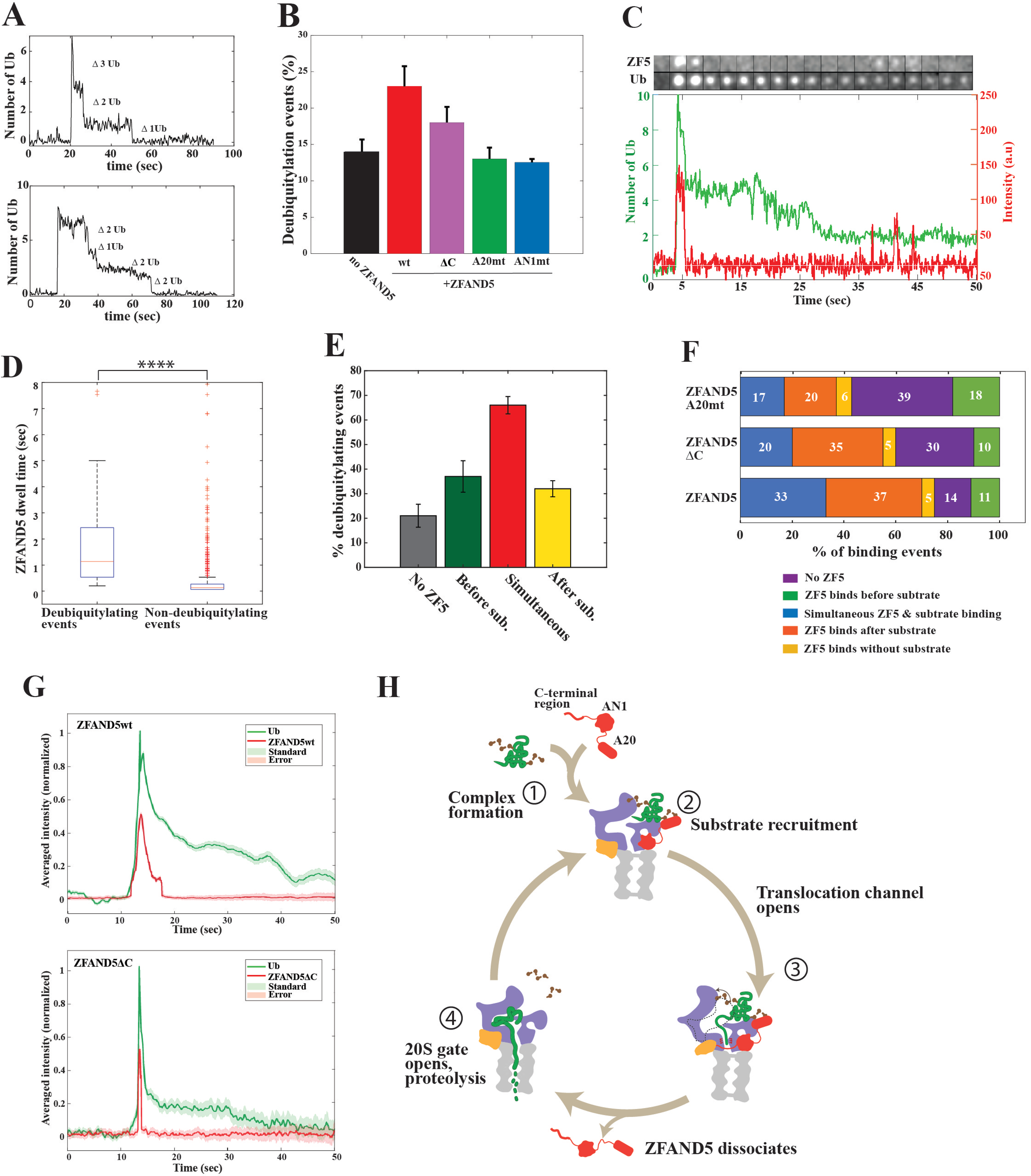
ZFAND5 ferries Ub conjugates to the proteasome and promotes their deubiquitylation and proteolysis. (A) Examples of single-molecule traces exhibiting processive deubiquitylation. Shown are the number of Ub molecules lost with each deubiquitylation step. (B) Effects of ZFAND5 and its mutants on the fraction of all substrate-proteasome encounters leading to processive deubiquitylation. Substrates containing at least four Ubs were analyzed. (C) An example montage and the time trajectory from a single-molecule measurement of ZFAND5 and ubiquitylated securin interacting with immobilized 26S proteasomes and undergoing dissociation or deubiquitylation. ZFAND5 was labeled with a JF646 dye via a SNAP tag; Ub was labeled with Dy550. (D) Comparison of ZFAND5 dwell times on proteasomes when leading to processive deubiquitylation or not. “*” indicates the statistical significance. (E) The fraction of substrate binding events leading to processive deubiquitylation when ZFAND5 binds with, before or after the substrate. See methods for the definition of each category. (F) ZFAND5, but not its A20 mutant or C-terminal deletion, increases the frequency of simultaneous binding with substrate to the proteasome. The fraction of times that the substrate binds together with, before, or after ZFAND5 or its mutants. (G) The averaged single-molecule kinetics of ZFAND5 and ubiquitylated securin on proteasome, suggesting that ZFAND5 dissociates from proteasomes before securin undergoes deubiquitylation. Single-molecule events recorded in the presence of ZFAND5 (N=440) or ZFAND5ΔC (N=97) were aligned by the moment of securin binding to proteasome, and the average fluorescence intensities among these events were plotted for the Ub and ZFAND5 channels. (H) ZFAND5 stimulates Ub conjugate degradation through a novel multistep reaction cycle 1) Formation of ZFAND5-substrate complex. 2) Its association with the proteasome RP. 3) RP assumes an open-channel conformation (Z+**_D_**) that favors deubiquitylation and degradation. 4) ZFAND5 dissociation leads to CP-gate opening, alignment with ATPase channel and substrate translocation into CP.

The increased probability of a bound substrate being degraded is not a simple consequence of its longer dwell time on the proteasome. At least for substrates with more than four Ubs, their dwell time on the proteasome is significantly longer than the average latency period of ∼2 seconds before the first deubiquitylation event, and therefore dwell time is not limiting for their degradation ^12^. Furthermore, the addition of ZFAND5 or its mutants does not significantly alter the interval from the initial substrate-proteasome encounter to the first or second deubiquitylation events (Fig. S14A), suggesting a dwell-time-independent mechanism enhances the degradation of these substrates that already have a high affinity for the proteasome.

Although increasing the likelihood of a substrate becoming committed to proteolysis, ZFAND5 does not boost the speed of translocation. We aligned the traces showing processive deubiquitylation by the moment of substrate binding and then calculated the average Ub intensity among these traces at each time point. We used the decay rate of the averaged trace as a measure of the speed of translocation ^12,31^. Neither ZFAND5 nor its mutants significantly affected the translocation speed of these substrates (Fig. S14B). This lack of effect on substrate translocation is consistent with our finding by cryo-EM that ZFAND5 is absent from the open-gated (i.e. translocation competent) **Z‒** states.

### The long-binding mode of ZFAND5 favors substrate degradation

The ZFAND5-induced **Z+_D_** state features a wide-open entrance to the ATPase translocation channel (Fig. 2B), which is likely to facilitate the capture of the substrate by the ATPases. To examine whether **Z+_D_** may underlie the increased substrate degradation by ZFAND5, we simultaneously recorded Ub’s and ZFAND5’s fluorescence signals in a dual-color single-molecule measurement (Fig. 7C). A variety of events were detected, which we first classified according to whether they were linked to processive deubiquitylation. ZFAND5 interacted with the proteasome transiently even in the presence of substrate. The median dwell time of ZFAND5 on the proteasome was 1.3s, if ZFAND5 binding occurred within 2s of the substrate binding and was associated with processive deubiquitylation (Fig. 7C and 7D). However, its dwell time was much shorter when ZFAND5’s binding events were not associated with substrate deubiquitylation. Because the **Z+_D_** state with its characteristic CTR-Rpn1-Rpt1 interactions appears responsible for the long-binding mode of ZFAND5, which is associated with substrate deubiquitylation, the novel **Z+_D_** state is very likely to be a critical step in ZFAND5’s stimulation of proteasomal degradation.

### Simultaneous ZFAND5 and substrate binding to the proteasome maximally stimulates proteolysis

We then compared the fractions of deubiquitylation events that occurred when ZFAND5 bound shortly before, together with, or shortly after the ubiquitylated substrate (Fig. 7E). Simultaneous binding (within one frame) of ZFAND5 and the substrate to the proteasome led to processive deubiquitylation and translocation in 65% of the cases, which is much higher than the likelihood of translocation (19%) seen without ZFAND5. When ZFAND5 binding preceded or followed substrate binding, it also increased the likelihood of degradation, but to a smaller extent.

The accelerated degradation of ubiquitylated substrates by ZFAND requires its A20 domain which interacts with Ub (Fig. 3C). To clarify further the function of A20, we analyzed their effects on the relative timing of the substrate binding to the 26S. Mutating the A20 domain or truncating the CTR decreased the probability of simultaneous binding of ZFAND5 and substrate to the proteasome, which is consistent with the importance of these domains in mediating substrate-ZFAND5 interaction (Fig. 7F). Thus, the timing of ZFAND5 and substrate binding to proteasome, which is mediated by the A20 and the CTR, determines the magnitude of the stimulation of conjugate degradation.

### ZFAND5 dissociates from the proteasome before substrate deubiquitylation and degradation

When the events where ZFAND5 and substrate bound simultaneously were aligned by the moment of initial substrate binding, the averaged signal from the ZFAND5 channel revealed that it participated only in the early stages of the degradation process. After substrate translocation started, ZFAND5 rapidly dissociated from the proteasome (Fig. 7G), as was also suggested by our structural analysis (Fig. 1). Similar rapid dissociation of ZFAND5 was evident when we analyzed only the deubiquitylation events (Fig. S14C). The loss of ZFAND5’s CTR significantly reduced its dwell time and substrate processing after the simultaneous binding, which further supports our conclusion that ZFAND5’s long-binding mode on the proteasome requires the CTR-dependent **Z+_D_** state and is critical in the acceleration of proteolysis. Together, these results indicate that ZFAND5 increases the likelihood that a bound substrate is committed to proteolysis through a series of coordinated events (Fig. 7H). To achieve this most effectively, ZFAND binds to the proteasome together with the ubiquitylated substrate. ZFAND5 then quickly dissociates, which transitions the proteasome to translocation-competent Z-states, where the gate in the CP is open and aligned with the ATPases (Fig. 7H).

## Discussion

### Molecular Mechanisms of Proteasome Activation

Recent studies have advanced markedly our understanding of proteasome function and have defined a series of conformational changes that catalyze the degradation of ubiquitylated proteins. This study differs in exploring for the first time the molecular mechanisms for the acceleration of Ub conjugate degradation induced by an important regulator of protein degradation, ZFAND5. Through a combination of Cryo-EM and single-molecule kinetic analysis, we have elucidated a coherent sequence of events underlying the ZFAND5-induced enhancement of 26S proteasome function, which hopefully will illuminate the mechanisms for proteasome activation by other physiological factors.

As summarized in Fig. 7H, (1) ZFAND5 stimulates most effectively when it initially associates with a ubiquitylated substrate through its A20 domain and its CTR and then together with the substrate binds to the proteasome. (2) On the proteasome, ZFAND5’s AN1 domain docks onto the Rpt5 ATPase subunit, which may release its CTR from the substrate, enabling it to engage with Rpt1 and Rpn1 on its Z site. (3) The binding of the ZFAND5’s CTR to Rpt1 and Rpn1 induces a novel conformation (**Z+_D_**) with a greatly enlarged ATPase channel, which should strongly facilitate substrate capture by the ATPases and the subsequent translocation. These interactions of the CTR with Rpt1 and the **Z** region of Rpn1 are of special regulatory importance, since a 19-residue CTR peptide can by itself enhance proteasomal hydrolysis of peptides and ubiquitylated proteins (see below). (4) ZFAND5 interacts only transiently with the proteasome and the substrate, and then rapidly dissociates, perhaps through an ATPase-driven step. (5) Importantly, its dissociation from the RP then converts the proteasome into a translocation-competent (**Z**-) state in which the CP gate that controls substrate entry is open and is aligned with the translocation channel in the RP. (6) Although the rate of substrate translocation does not appear to be accelerated, the likelihood that a bound Ub conjugate becomes committed to degradation is markedly increased through this sequence of events.

It was initially puzzling to find that ZFAND5 could stimulate proteasomal degradation even though it was only present in the translocation-incompetent (i.e. closed-gate) states. However, our dual-color single-molecule experiment (Fig. 7) showed that ZFAND5 functions primarily at the initial steps in this sequence, and that its dissociation precedes deubiquitylation and transition to translocation-competent **Z‒** states, some of which had only been observed previously in substrate-engaged 26S structures ^12^. This rapid dissociation indicates that ZFAND5 does not function as an “alternative receptor” that holds the substrate on the RP in a distinct position ^20^, but instead serves to trigger critical, structural changes. The transition into these **Z‒** states requires and likely follows the open-entry **Z+_D_** state, because the ZFAND5’s CTR is essential for stimulating the 26S’s proteolytic activities, but not for the initial interaction with the proteasome (Fig. 3, 5 and 7). Although ZFAND5 stimulates proteolysis most effectively, when it binds to the proteasome together with the ubiquitylated substrate, there exists a short time window of approximately ± 2 sec after the binding of either the substrate or of ZFAND5 during which a stimulatory effect of ZFAND5 is manifest. Presumably this means that the ubiquitylated substrate can utilize these ZFAND5-induced transitions even if not initially bound to ZFAND5. This conclusion is also supported by our observations with the 19 residue CTR peptide, which lacks the Ub conjugate-binding A20 domain, but still stimulates proteasomal activities.

Because ZFAND5’s capacity to bind ubiquitylated substrates and promote their degradation requires its A20 domain, it was quite surprising to find that the synthetic CTR peptide could alone stimulate degradation of both short peptides and Ub conjugates (although less effectively than the complete molecule). These observations further suggest that the CTR-dependent **Z+_D_** state is critical in mediating ZFAND5’s stimulatory effects, and that the interaction with the substrate is not absolutely necessary for activation. The requirement for much higher concentrations of this peptide than for ZFAND5 is probably because the docking of the AN1 domain on Rpt5 enhances ZFAND5’s affinity for the proteasome and increases the ability of the CTR to associate with the **Z** regulatory site on Rpn1. It is noteworthy that the CTR peptide and ZFAND5 seem to affect the multiphase kinetics of degradation of a Ub conjugate rather differently. The peptide was most effective (which means that the A20 domain is dispensable) in increasing conjugate degradation during the outset of phase I of the reaction (Fig. 5B), where substrates bearing high numbers of Ub are preferentially degraded. However, the peptide had no effect, in phase II, where ZFAND5 primarily enhances degradation of substrates that have low numbers of Ub and interact weakly with the proteasome, and therefore the A20 and AN1 domains are essential.

In its ability to interact simultaneously with the proteasome and the ubiquitylated substrate, and thus to function in substrate recruitment, ZFAND5 seems to resemble one of the UBL-UBA “shuttling factors” (i.e. Rad23, Dsk2, Ddi2, ubiquilins), although there are no sequence similarities between them. *In vivo* the shuttling factors deliver ubiquitylated substrates to the 26S proteasome and have been assumed to enhance their hydrolysis ^11^, although these properties have not been directly demonstrated in vitro. The binding of a protein bearing a UBL domain to the proteasome does stimulate 26S peptide hydrolysis (CP gate opening), but unlike ZFAND5, these UBL-proteins by themselves do not increase Ub-conjugate degradation or ATP hydrolysis ^33,34^. On the proteasome, the UBL -domains activate peptide hydrolysis by binding to Rpn1 at its T1 and T2 sites. Although these sites are distinct from the **Z** site, it is possible that occupancy of these different regulatory regions on Rpn1 induce some similar structural changes leading to activation.

While ZFAND5 and other activators of the 26S proteasome (e.g. PKA or PKG) appear to enhance overall proteolysis in cells ^1,6^, they probably do not stimulate the degradation of all ubiquitylated cell proteins similarly. In our experiments using substrates of the N-end rule pathway and the UFD pathway, ZFAND5 expression increased their degradation, which depends on its A20, AN1 and CTR domains, similarly as in the purified reactions. Also, the degree of activation by ZFAND5 depends on the number of Ub on the substrate and perhaps on the pattern of ubiquitylation. Exactly how the A20 domain and the CTR of ZFAND5 interact with Ub conjugates is unclear especially since there are no recognizable Ub-binding elements within the CTR. Nevertheless, this association may also lead to the selection of substrates with certain Ub configurations. Also after binding to the 26S, some features of the polypeptide are likely to be important in substrate selection.

It is noteworthy that ZFAND5 affects multiple steps in the degradative process, and it is unlikely that any specific step in this reaction cycle by itself accounts for the more rapid degradation. ZFAND5’s capacity to enhance substrate association with the proteasome (i.e. prolong dwell time), to enlarge the entrance into the ATPase channel, and upon dissociation to induce CP gate-opening all seem likely to help stimulate proteolysis. Moreover, there are probably additional steps stimulated by ZFAND5. Our experiments with difficult-to-degrade substrates suggested that ZFAND5 may also increase their deubiquitylation by Rpn11, which can be a rate-limiting step for proteolysis. The present studies have examined conditions with the substrate excess over the proteasome; so possible effects of ZFAND5 on promoting substrate binding (K_on_) should have little impact, but could be important *in vivo*. It is also noteworthy that even in the absence of a substrate, ZFAND5 stimulates ATP hydrolysis by the proteasome ^1^. The molecular basis for the faster ATPase is an interesting question for future work; perhaps this process is activated when the RP assumes a more open conformation (**Z+_D_**) or with ZFAND5 dissociation and the alignment of the ATPases with the open gate in the CP. In any case, greater ATPase activity should also enhance proteolysis, since the rate of Ub conjugate degradation by proteasomes is proportional to their rate of ATP consumption ^35^. Finally, the entire degradative process must be more efficient with ZFAND5 present, since it greatly decreases the frequency that a substrate binds to the proteasome but fails to be degraded.

### Biological implications and potential applications of these findings

It is widely assumed that 26S proteasomes are identical in different tissues, but the expression of ZFAND5 normally in heart and brain and its induction in atrophying muscles strongly suggests that in these tissues, proteasomes are more active and consequently the destruction of ubiquitylated proteins is more efficient than in other cell types. Cryo-EM tomography of hippocampal neurons and green algae has led to the conclusion that the great majority of proteasomes in eukaryotes are normally in an inactive conformation ^36,37^. However, this conclusion cannot apply to cells where proteasomes are activated (heart or brain, atrophying muscles or cells upon heat shock or stimulation by certain hormones)^1,5,38^. Our use of covalent probes of the CP’s active sites has proven a valuable tool to monitor the 26S activation *in vitro* and in cells ^39^ and should enable other studies of proteasome activation. Future studies using a similar combination of Cryo-EM, rigorous biochemical characterization, and single-molecule kinetic analysis will be necessary to clarify the possible similarities and differences between ZFAND5’s actions and these other 26S activators.

In muscle during fasting or atrophy, when overall rates of proteolysis rise, the levels of ubiquitylated substrates also are increased ^40^; thus proteasomal function must limit the rate of degradation, and 26S activation would seem important for the rapid loss of muscle mass. Muscle proteins, primarily the components of the myofibrillar proteins, are the major protein reservoir in mammals, and its mobilization in fasting and disease provides key precursors for gluconeogenesis and energy production ^41^. In these catabolic states, there is a selective destruction of these normally very stable myofibrillar components by the UPS. Thus, there is a clear rationale for the enhanced proteasomal activities, and perhaps specifically for an increased capacity to digest difficult-to-degrade proteins. The possible physiological advantage of normally maintaining proteasomes in an activated state in heart or brain is an intriguing question; certainly these are tissues where protein quality control appears of major medical importance.

The important role of ZFAND5 in the accelerated proteolysis driving muscle atrophy implies that inhibitors of its actions may have therapeutic applications in combatting the debilitating loss of muscle mass seen with nerve injury, inactivity and many systemic diseases (e.g. cancer cachexia, cardiac and renal failure, excess glucocorticoids). Our identification of the critical interactions between ZFAND5 and the proteasome, especially those between its CTR and Rpt1 and Rpn1’s Z-site should facilitate the development of small molecule inhibitors of ZFAND5-dependent muscle wasting. Conversely, for the various proteotoxic diseases, where the accumulation of misfolded, aggregation-prone proteins impairs proteasome function ^42^, overexpression of ZFAND5 or small molecules that mimic its ability to activate proteasomes would be a useful therapeutic strategy. The ability of the 19 residue C-terminal peptide to enhance by itself Ub conjugate degradation strongly suggests that its interactions with the RP should be an attractive target for pharmacological development of proteasome activators.

## Limitations of the study

Here, we presented structural, functional and kinetic measurements to elucidate the mechanism of proteasome activation by its cofactor ZFAND5 which has been implicated in several important biological processes. However, it is still unclear whether ZFAND5 affects all proteasome substrates equally or imposes a selectivity which may be important for understanding its biological functions. Technically, combining substrates with ZFAND5 in structural study may provide further insights into the activation process. Moreover, the cell contains a number of factors and modifications that are able to activate the proteasome. Further studies should address these questions and uncover the complex regulation of proteasome activity and its role in protein homeostatsis.

## Acknowledgements

We are grateful for funding from the National Institute of General Medical Sciences (R01 GM134064-01 to YL; R01 GM051923-20 to ALG), the Cure Alzheimer’s Fund, the Muscular Dystrophy Association (to ALG), the Edward Mallinckrodt Jr. Foundation Award to YL, NIH R01GM074830 and R35GM145249 to LH, National Natural Science Foundation of China (NNSFC12090054 to QO), and want to thank Ms Amelia Gould for valuable assistance in preparation of this manuscript. We want to thank Jack (Youdong) Mao, Weili Wang and Xuemei Li at Peking University for assistance in cryo-EM technology. The cryo-EM data were collected from the Electron Microscopy Laboratory and Cryo-EM Platform at Peking University. Data processing was supported by High-Performance Computing Platform at Peking University.

## Author contributions

D. L., A. L. G and Y. L. conceived and designed the project. D. L. purified proteins and performed biochemical analysis. Y. Z. performed cryo-EM analysis and single-particle reconstruction of different proteasome states with ZFAND5. L. C. performed single-molecule fluorescence analysis. D. L. and S. C. performed live-cell analysis. L. H. and X. W. performed mass spec analysis. All authors participated in manuscript preparation.

## Data availability

Cryo-EM maps haven been deposited in the Electron Microscopy Data Bank (EMDB) under accession codes EMD-14201 (Z+_A_), EMD-14202 (Z+_B_), EMD-14203(Z+_C_), EMD-14204 (Z+_D_), EMD-14205 (Z+_E_), EMD-14209 (Z‒_A_), EMD-14210 (Z‒_B_), EMD-14211 (Z‒_C_). Coordinates are available from the RCSB Protein Data Bank under accession codes 7QXN (Z+_A_), 7QXP (Z+_B_), 7QXU (Z+_C_), 7QXW (Z+_D_), 7QXX (Z+_E_), 7QY7 (Z‒_A_), 7QYA (Z‒_B_), 7QYB (Z‒_C_).

## STAR★METHODS

### KEY RESOURCES TABLE

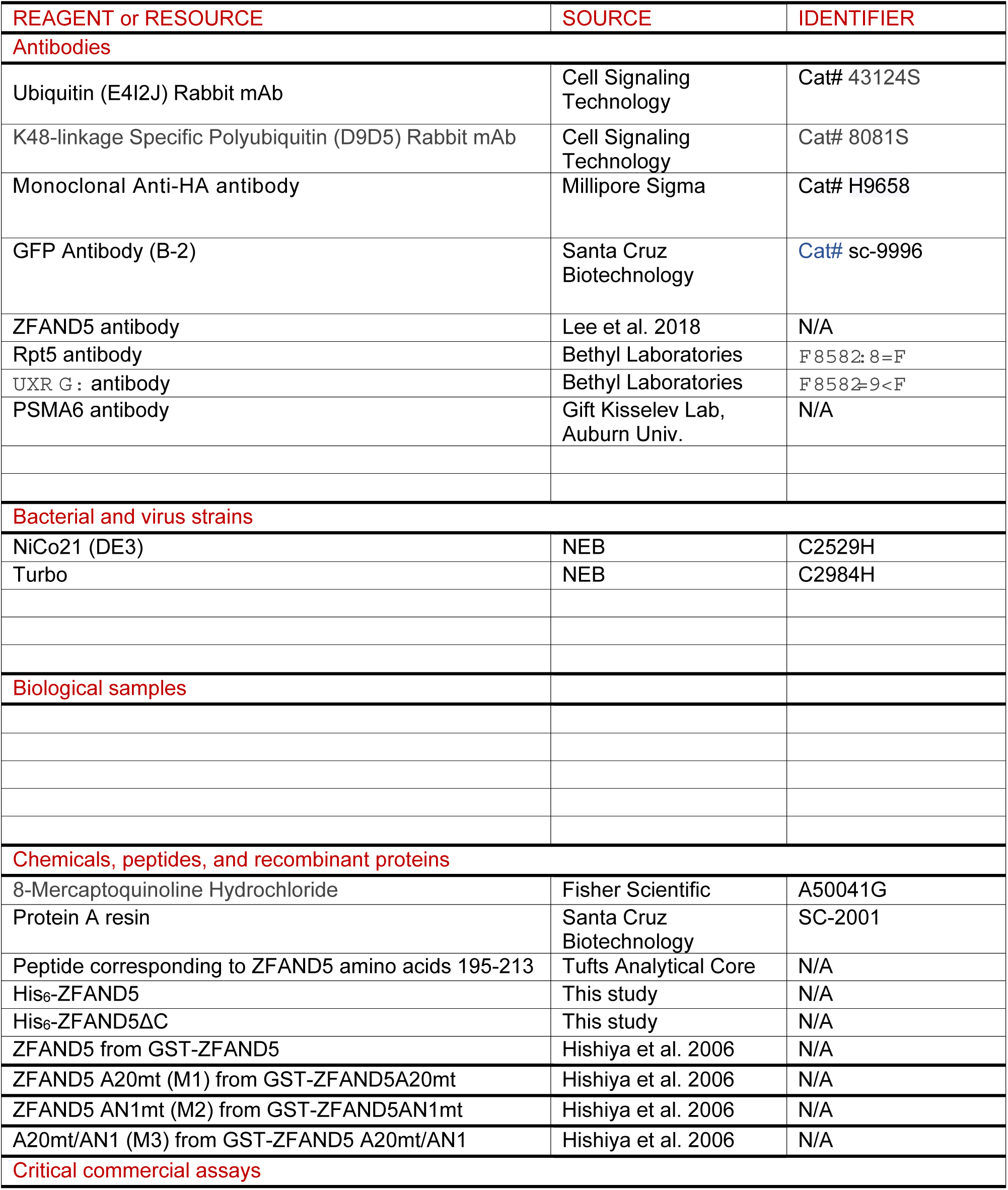
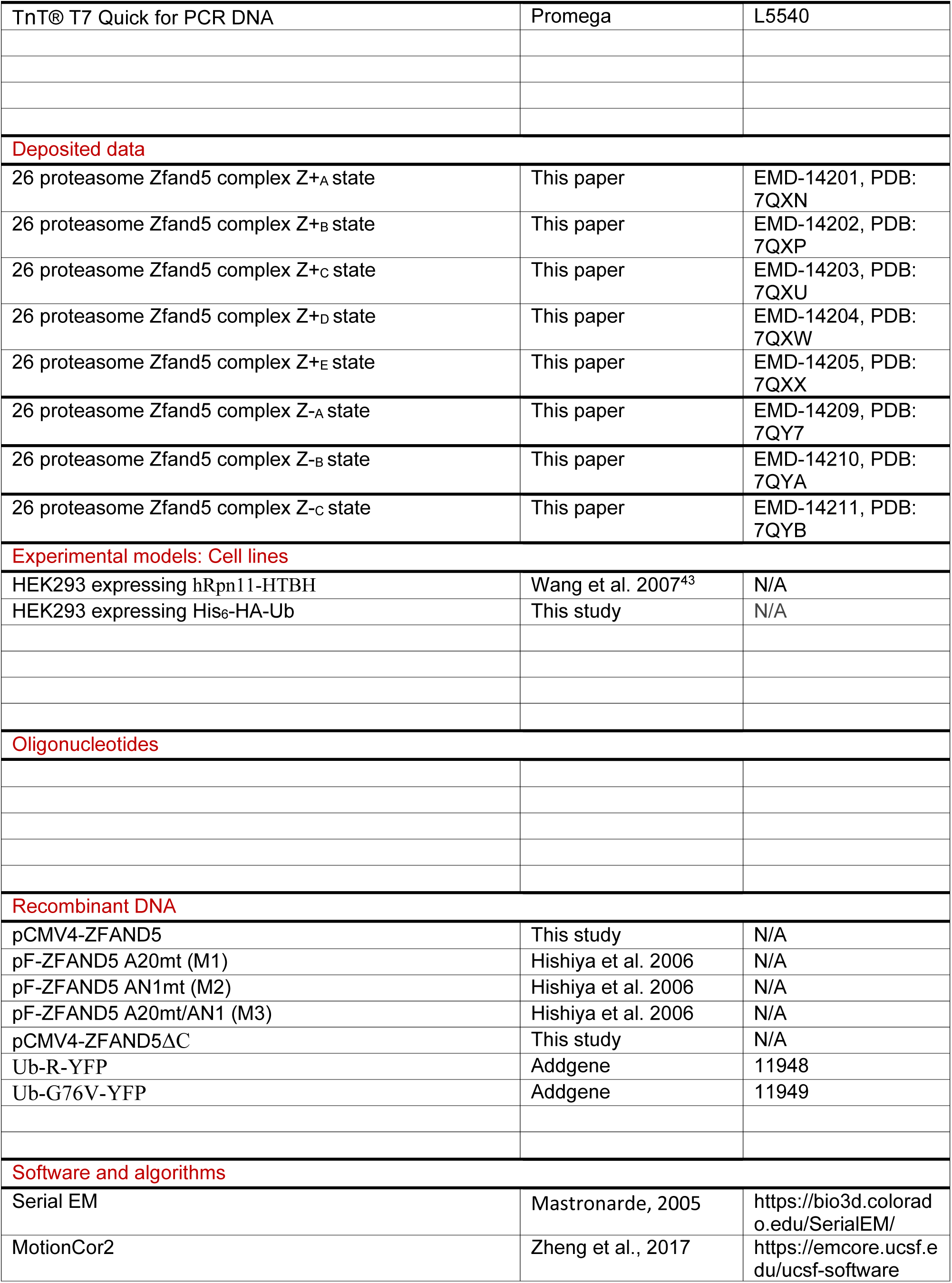
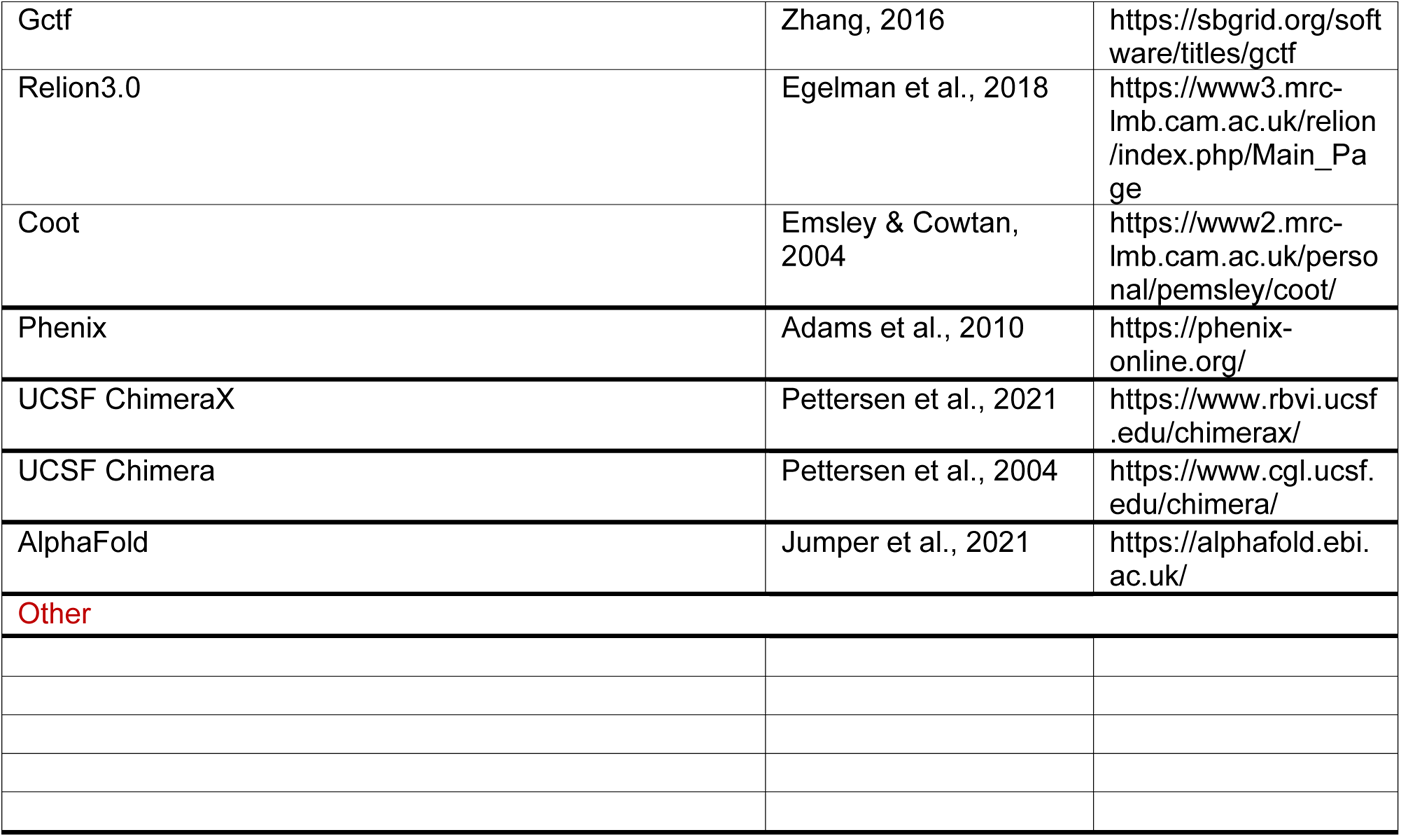

### RESOURCE AVAILABILITY

#### Lead contact

Ying Lu. Ph. D. Dept. of Systems Biology, Harvard Medical School Alfred Goldberg, Ph. D. Dept. of Cell Biology, Harvard Medical School

#### Materials availability

This study did not generate new unique reagents.

#### Data and code availability

Cryo-EM maps haven been deposited in the Electron Microscopy Data Bank (EMDB) under accession codes EMD-14201 (Z+A), EMD-14202 (Z+B), EMD-14203(Z+C), EMD-14204 (Z+D), EMD-14205 (Z+E), EMD-14209 (Z‒A), EMD-14210 (Z‒B), EMD-14211 (Z‒C). Coordinates are available from the RCSB Protein Data Bank under accession codes 7QXN (Z+A), 7QXP (Z+B), 7QXU (Z+C), 7QXW (Z+D), 7QXX (Z+E), 7QY7 (Z‒A), 7QYA (Z‒B), 7QYB (Z‒C).

### EXPERIMENTAL MODEL AND SUBJECT DETAILS

### METHOD DETAILS

#### Proteasome expression and purification

Human proteasomes were purified through affinity chromatography on a large scale from a stable HEK293 cell line with RPN11-HTBH (hexahistidine, TEV cleavage site, biotin, and hexahistidine) (Wang et al., 2007). The harvested cells were homogenized with a Dounce homogenizer (type B pestle or equivalent tight one) in a lysis buffer (50 mM NaH2PO4, pH 7.5, 100 mM NaCl, 5 mM MgCl2, 0.5% NP-40, 5 mM ATP, 1 mM DTT) containing protease inhibitor cocktail (Roche, Germany). The lysates were cleared, and incubated with the NeutrAvidin agarose beads (Thermo Fisher Scientific, MA, USA) overnight at 4°C. The beads were washed by excess lysis buffer and followed by a wash with TEB buffer (50 mM Tris-HCl, PH 7.5) containing 10% glycerol and 1 mM ATP-MgCl2. The proteasome holoenzymes were eluted from the beads through cleavage by TEV protease (Invitrogen, CA, USA). The doubly capped proteasome was further purified by gel filtration on a Superose 6 10/300 GL column (GE Healthcare, PA, USA) at a flow rate of 0.15 ml/min in the running buffer (30 mM Hepes pH 7.5, 60 mM NaCl, 1 mM MgCl2, 10% Glycerol, 0.5 mM DTT, 0.8 mM ATP). The gel-filtration fractions were concentrated to about 2 mg/ml. Measurement of peptidase activity of purified 26S complexes was performed as previously described (Lee et al., 2018).

#### Cryo-EM Data collection

Immediately before cryo-EM sample preparation, the proteasome sample was mixed with ZFAND5 with a molar ratio about 1:100 and buffer-exchanged into 50 mM Tris-HCl pH 7.5, 1 mM MgCl2, 1 mM ATP, and 0.005% NP-40 using a 7-kDa Zeba column. Cryo-EM sample grids were prepared using the FEI Vitrobot Mark IV (Thermo Fisher Scientific, MA, USA). C-flat grids (R1.2/1.3; 400 Mesh, Protochips, CA, USA) were glow-discharged before a 2.5-μl drop of proteasome-ZFAND5 mixed solution was applied to the grids in an environment-controlled chamber with 100% humidity and temperature fixed at 4 °C. After 3 s of blotting, the grid was plunged into liquid ethane and then transferred into liquid nitrogen. The cryo-grids were imaged using a FEI Titan Krios microscope (Thermo Fisher Scientific, MA, USA), equipped with an Autoloader and operating at an acceleration voltage of 300 kV at a nominal magnification of 105,000 times. Cryo-EM movie data were collected using serialEM software on a Gatan K2 Summit direct detector camera in a super-resolution counting mode, with 10 s of total exposure time and 250 ms per frame. Each exposure resulted in a movie of 40 frames with an accumulated dose of 46.6 electrons/Å^2^. The calibrated physical pixel size and the super-resolution pixel size were 1.37 and 0.685 Å per pixel, respectively. The defocus was prescribed in the range from - 0.8 to - 2.5 μm. A total of 8653 movies in super-resolution mode were collected for data analysis.

#### Cryo-EM data processing and reconstruction

All frames of the raw movies were first corrected for their gain using a gain reference recorded within 3 days of the acquired movie, after which they were shifted and summed to generate a single micrograph that was corrected for overall drift using the MotionCor2 (Zheng et al., 2017). Each drift-corrected micrograph was used for the determination of the actual defocus of the micrograph using Gctf program (Zhang, 2016). Particle picking was done using DeepEM program (Zhu et al., 2017) and 501,131 single particle images were picked for further analysis. The density map of previously published SA (Zhu et al., 2018) state was low-passed filtered to 60 Å and been used as the initial model.

All the 2D/3D classification and auto-refinement were done in Relion3.0 (Egelman et al., 2018). The first round 2D/3D classification was done at a pixel size of 2.74 Å and classified the data to two parts, double-capped proteasome with 272,785 particles and single-capped proteasome with 137,130 particles. Then we re-centered particles on RP-CP sub-complexes and converted each double-capped particle to two pseudo single-cap particles by shifting the center to each side (Dong et al., 2019; Zhu et al., 2018). We re-extracted 682,700 pseudo single-cap particles. The second round 2D/3D classification was also done at a pixel size of 2.74 Å and classified the data to CP-gate closed states with 35,4682 pseudo single-cap particles and CP-gate open states with 199,312 pseudo single-cap particles. All the particles in the CP-gate closed states were merged to one class and all the particles in CP-gate open states were merged to another class. Both classes run auto-refinement at the pixel size of 1.37 Å. In the third round, we did focus 3D classification at the pixel size of 1.37 Å with a RP-mask and only did local angular search with angular range of 7.5 degree based on the orientations determined by the auto-refinement in last round. Five different conformational states named state Z+A, Z+B, Z+C, Z+D and Z+E were determined from the CP-gate closed class with particle numbers 80,224, 90,097, 29,419, 46,247 and 28,928 respectively. Three conformations states named Z-A, Z-B and Z-C were determined from CP-gate open class with particle numbers 34,363, 59,461 and 34,075 respectively. After auto-refinement and CTF refinement, the final overall resolutions for Z+A, Z+B, Z+C, Z+D, Z+E, Z-A, Z-B and Z-C are 3.7Å, 3.6Å, 4.3Å, 4.1Å, 4.4Å, 4.7Å, 4.1Å and 4.8Å respectively, measured by gold-standard FSC at 0.143-cutoff on two half maps refined separately. Prior to visualization, all density maps were sharpened by applying a negative B-factor calculated by Relion post-process (Egelman et al., 2018). Local resolution variations were estimated using Relion local resolution estimation (Egelman et al., 2018).

#### Atomic model building and refinement

The initial atomic models for 26S proteasome were based on previously published substrate-engaged human 26S structures (Dong et al., 2019) and then manually improved the main-chain and sidechain fitting in Coot (Emsley & Cowtan, 2004) to generate the starting coordinate files. The initial atomic model for ZFAND5 was based on the NMR structure with PDB ID:1WFL and AlphaFold predicted structure (Jumper et al., 2021). To fit the model to the reconstructed density map, we first conducted rigid-body fitting of the segments of the model in Chimera (Pettersen et al., 2004), after which the fit was improved manually in Coot (Emsley & Cowtan, 2004). Finally, each refinement of the atomic model was carried out in real space with program Phenix.real_space refine (Adams et al., 2010), with secondary structure and geometry restrains to prevent overfitting.

#### Structural analysis and visualization

Structural comparison and visualization were conducted in UCSF ChimeraX and Chimera. All figures of the structures were plotted in UCSF ChimeraX (Pettersen et al., 2021) and Chimera(Pettersen et al., 2004).

#### DSSO Cross-linking of Affinity Purified 26S Proteasome-ZFAND5 Complex

The stable 293 cells expressing Rpn11-HTBH were grown to ∼90% confluence in a DMEM medium containing 10% FBS and 1% Pen/Strep as previously described (Wang et al., 2007). The cells were pelleted and washed with PBS and then lysed in a native lysis buffer [100 mM sodium chloride, 50 mM sodium phosphate, 10% glycerol, 1 mM DTT, 5 mM MgCl2, 1 mM ATP, 1× protease inhibiter (Roche), 1× phosphatase inhibitor, and 0.5% NP-40 (pH 7.5)]. The lysates were centrifuged at 13,000 rpm for 15 min to remove cell debris, and the supernatant was incubated with streptavidin resin 2 hours at 4 °C. The streptavidin beads were then washed with 50 bed volumes of the lysis buffer, followed by a final wash with 20 bed volumes of crosslinking buffer (150 mM sodium chloride, 25 mM sodium phosphate, 5% glycerol, 5 mM MgCl2, 1 mM ATP). Bound proteasomes were incubated with equal volume (bed volume) of 70µM ZFAND5 at 37°C for 30 min, then DSSO was added to the mixture at final concentration of 0.5 mM and incubate for 1 h at 37°C. After quenching the cross-linking reaction, the proteins were reduced/alkylated and digested with LysC/trypsin. Briefly, proteins were digested in 8 m urea buffer using LysC for 4 h at 37°C, followed by trypsin digestion at 37°C overnight after diluting urea concentration to <1.5 M. The resulting peptide mixtures were extracted and desalted before MS analyses.

#### Identification of DSSO Cross-Linked Peptides by LC MS^n^

Peptide digests were analyzed by LC MS^n^ using an UltiMate 3000 RSLC coupled with an Orbitrap Fusion Lumos mass spectrometer similarly as described (C. Yu et al., 2019). Samples were loaded onto a 50 cm x 75 μm Acclaim PepMap C18 column and separated over a 240 min gradient of 4% to 25% acetonitrile at a flow rate of 300 nL/min. The top 4 data-dependent MS^3^ acquisition method was used for the identification of DSSO cross-linked peptides. Ions with charge of 4+ to 8+ in the MS^1^ scan were selected for MS^2^ analysis. The top 4 most abundant fragment ions in MS^2^ scan were further fragmented by CID with a collision energy of 35%. Raw data were extracted by MSConvert and MS^3^ spectra were subjected to Protein Prospector (v.6.2.13) for database searching using Batch-Tag against SwissProt.2019. 04. 08 random concatenated database. The mass tolerances were set as ±20 ppm for parent ions and 0.6 Da for fragment ions. Trypsin was set as the enzyme with three maximum missed cleavages allowed. Cysteine carbamidomethylation was selected as fix modification. A maximum of three variable modifications were also allowed, including methionine oxidation, N-terminal protein acetylation, and N-terminal conversion of glutamine to pyroglutamic acid. Three defined DSSO cross-linked modification on uncleaved lysines, including alkene (C3H2O, +54 Da), thiol (C3H2SO, +86 Da) and sulfenic acid (C3H4O2S, +104 Da) were also selected as variable modifications. Search results were integrated via in-house software xl-Tools to identify DSSO cross-linked peptides by the integration of MS^1^, MS^2^ and MS^3^ data.

#### Purification of recombinant proteins

Recombinant ZFAND5 wild type and mutants containing point mutations in Zn-finger domains (ZFAND5A20mt or ZFAND5AN1mt) were purified as previously described(Lee et al., 2018). Bacterial expression plasmids encoding His6-ZFAND5ΔC, His6-SNAP-ZFAND5wt, His6-SNAP-ZFAND5ΔC, His6-HA-cycB-cpGFP, His6-HA-cycB-mNeon Green or His6-HA-cycB-EGFP were generated. All proteins containing hexahistidine-tag (His6), including E2 UbcH10, ubiquitin and Securin, were purified with Ni-NTA resin (Qiagen).

Recombinant Anaphase Promoting Complex (APC/C) and His6-Cdh1 were purified from insect cells as previously described (Lu et al., 2015).

#### Labeling proteasomes after purification and in cells with the activity-based probe

26S proteasomes (2nM) were incubated in the reaction buffer (50 mM Tris-HCl (pH 7.5), 100 mM KCl, 5 mM MgCl2, 1 mM ATP, 1 mM DTT) and MVB003 (500 nM) (gift from Herman Overkleeft) at 37°C with or without ZFAND5 (500 nM). To label proteasomes in atrophying C2C12 myotubes, cells pretreated with dexamethasone (50mM) for 1d were incubated with MVB003 (500 nM) for 1hr. Samples were resolved in SDS-PAGE and scanned with an AI600 RGB camera with λex 520nm and λem 593nm. Fluorescence of the bands was quantified with the IQTL software package from GE Healthcare Sciences.

#### Degradation assay with non-ubiquitylated ZFAND5 and 26S proteasome

His6-ZFAND5 was labeled during expression with 35S-methionine using the kit for coupled transcription/translation system (Promega) and purified with Ni-NTA resin. Purified His-ZFAND5 was incubated with 26S proteasomes up to 60min, and the reactions were stopped by addition of TCA at the indicated time points. Hydrolysis of ^35^S-His6-ZFAND5 to TCA-soluble peptides was measured, and the radioactivity in acid soluble and insoluble fractions are presented as the mean ± SD of three replicates.

#### In vitro ubiquitylation

Ubiquitylation of substrate was performed as described previously (Fang et al., 2022; Lu et al., 2015). Briefly, polyubiquitylation reactions were performed with 2μM of His6-HA-cycB-cpGFP, His6-HA-cycB-mNeonGreen or His6-HA-cycB-EGFP were by APC/C complex (50nM), 100 nM E1, 2μM UbcH10, 2 mg/ml bovine serum albumin (BSA), 10mM creatine phosphate, 0.1 mg/ml creatine kinase, 10μM ubiquitin in UBAB buffer (25mM Tris-HCl pH 7.5, 50mM NaCl and 10mM MgCl2) at room temperature for 4h.

#### Degradation of ubiquitylated substrates

The reaction mixture containing ubiquitylated substrates (120-200nM), 26S proteasomes (2-5nM) and ZFAND5 (0.03-5μM) was incubated in UBAB buffer at 35°C. Degradation of substrate was detected by measurement of fluorescence of cpGFP or EGFP (λex, 470 nm; λem, 510 nm) in 1min interval for 60min. The background level of fluorescent signal from the reaction buffer alone for each time point was subtracted from all the signals with the substrate. The fluorescent signals during degradation reaction were also normalized to small changes in fluorescence from a substrate alone, and then signals relative to that at 0 min were plotted in the figures. Each figure shows the data from at least three independent experiments with 3-5 replicates for each condition. The reactions with ubiquitylated cycB-EGFP were alternatively resolved in SDS-PAGE, and the change in cycB level was probed by the antibody to HA (H9658, Millipore Sigma), ubiquitylation by antibodies to K48-linked chain (#8081, Cell Signaling Technology) or total Ub (#43124, Cell Signaling Technology), and EGFP by GFP antibody (SC-9996, Santa Cruz Biotechnology).

#### Coimmunoprecipitation assay

Purified 26S proteasomes were immobilized with the antibody to α6 crosslinked on proteinA resin (SC-2001, Santa Cruz Biotechnology). ZFAND5wt or ZFAND5ΔC was added and incubated for 1h at 4°C. After washing with 300mM NaCl, resin-bound proteins were resolved in SDS-PAGE and probed by Western blot with antibodies to α6 (A303-845A, Bethyl Laboratories), Rpt5 (A303-538A, Bethyl Laboratories) and ZFAND5 (Lee et al., 2018).

ZFAND5 was incubated with resin-bound 26S proteasomes for 30min on ice, and resin was washed with the buffer used for degradation assay to remove any free ZFAND5. CTR peptides were then added to ZFAND5-bound 26S and incubated for 30min on ice. Unbound proteins or peptides were washed off with the reaction buffer, and the levels of proteasome-bound ZFAND5 were determined by Western blot.

#### Native gel

Purified 26S proteasomes were incubated with or without ZFAND5 for 20min at 37°C and the complexes were resolved in 3-8% gradient native-PAGE. Migration of 26S complexes was detected with the antibody to Rpt5.

#### Phosphorylation of ubiquitylated substrate

PKA site (RRASV) at the N-terminus of His6-HA-cycB-cpGFP was radiolabeled with ^32^P-ATP, and radiolabeled substrate was then polyubiquitylated by APC/C. Degradation assay was performed as describe above and stopped at the indicated time points in the figure. The levels of ubiquitylated substrates were quantified by a phosphorimager (Typhoon5).

#### Single-molecule measurement of the kinetics of substrate processing by proteasome in the presence of ZFAND5

Recombinant ubiquitin was labeled with dylight550-maleimide at the N-terminus. The N-terminal His tag was then cleaved off by thrombin and was purified using anion exchange FPLC. Securin labeled with dy550-ubiquitin (Securin-Ub550) was prepared as described previously (Lu et al., 2015). Purified 26S proteasome and biotinylated MCP21 antibody were mixed to a final concentration of 20 nM and 12.5 nM respectively. The mixture was incubated at room temperature for 15 min then kept on ice until the experiment. For all imaging experiments, the temperature was set to 29°C ±2°C, unless indicated otherwise. The proteasome-antibody mix was loaded onto passivated slides coated with streptavidin and incubated for 3 min.

For the single-color imaging, unbound proteasome was washed off and replaced with imaging buffer containing diluted 5 nM ubiquitylation product and 500 nM purified ZFAND5 (WT, AN1mt, A20mt, or ΔC). Image acquisition was started immediately with <15 s delay. Time series were acquired at 200 ms per frame for 3 min.

For the dual-color imaging, 2 μM 649-SNAP-ZFAND5 or 2 μM 649-SNAP-ZFAND5ΔC was prepared by incubating 2 μM of purified SNAP-ZFAND5 with 3 μM SNAP-Surface 649 for 1 hour at room temperature and purified by desalting column. For the experiments, unbound proteasome was washed off and replaced with imaging buffer containing diluted 17 nM ubiquitylation product and 50 nM of either 649-SNAP-ZFAND5 or 649-SNAP-ZFAND5ΔC. Image acquisition was started immediately with <15 s delay. Time series were acquired at 50 ms per frame for 4000 frames (∼3 min). TetraSpeck™ Microspheres, 0.1 µm (T7279) were used to assist color channel registration.

We used a Nikon Ti TIRF microscope equipped with three laser lines of 488 nm (OBIS™ 1277611), 561 nm (Opto Engine MGL-FN-561-100mW, ∼1mW at the objective), and 638 nm (Lasever LSR635NL 150 mW, ∼0.2mW at the objective), a Nikon Plan Apo λ 100X/1.45 Oil objective, and a Pco.edge 4.2 LT HQ camera.

#### Single-molecule measurement of Ub conjugates binding to ZFAND5

Biotinylated zfand5 was prepared by incubating 2 μM SNAP-ZFAND5 with 1 μM SNAP-Biotin^®^ (S9110S) in UBAB for 1 hour at room temperature. The 2uM biotinylated ZFAND5 was loaded onto passivated slides coated with streptavidin and incubated for 5 min. Unbound ZFAND5 was washed off with imaging buffer. Securin-Ub550 diluted in imaging buffer was flowed in at a given concentration and image acquisition was started immediately with <15 s delay. Time series were acquired at 30 ms per frame for 1000 frames (30 seconds).

#### Analysis of single-molecule data: dwell time, deubiquitylation/translocation kinetics, and dual color processing

Image processing was performed as described previously (Lu et al., 2015). In short, image sequences were corrected for stage drift, a custom spot-detection algorithm with <5% false-positive rate identified spots, and the intensity of each spot was obtained by fitting with a two-dimensional Gaussian function. The signal was converted to the copy number of ubiquitin on each substrate molecule by normalizing with the intensity value of a single ubiquitin-550 obtained in the photobleaching calibration step as described previously(Lu et al., 2015). The custom-built algorithm (Lu et al., 2015) was used to measure the duration of substrate-binding events. In the dwell time measurement, we did not differentiate whether or not a binding event exhibited processing deubiquitylation.

The measurement of substrate translocation kinetic was performed as previously described(Lu et al., 2015). Briefly, a custom-built algorithm was used to align the start of all single-molecule traces showing processive deubiquitylation, i.e. translocation, by the moment of substrate-proteasome interaction. The alignment was manually curated to remove false alignment. The substrate-proteasome interaction was identified by finding the first timepoint where the intensity is at least 80 percent of the maximum intensity within the manually determined start and end times. After alignment, the averaged translocation kinetics was calculated from the intensity average among all the traces at each time point.

For processing dual color imaging, the image sequence was parsed into the two channels. The Securin-Ub550 channel image sequence was processed in the same was as described above. The ZFAND5 sequence inherited the corrections for stage drift and the positions of the identified spots from the Ub550 channel image sequence processing. Traces were aligned by the moments of substrate-proteasome interaction, and this alignment was applied to both the ubiquitin and ZFAND5 channel.

#### Classification of single-molecule binding events

Single-molecule binding events involving both substrate and ZFAND5 were classified manually. Signals lasting only 1 frame (50ms) were not considered as binding events to reduce the influence of background fluctuation.

For the dual color events, the role of ZFAND5 was classified manually:

- The event was determined to have “no ZFAND5” if there was no ZFAND5 signal during or within 10 sec of either side of the Securin-Ub550 binding event.
- If ZFAND5 signal appeared within 10 seconds before the Securin binding event, the event was classified as “ZFAND5 binds before substrate”.
- The event was classified as “simultaneous ZFAND5 & substrate binding” if ZFAND5 signal appeared within one frame of the Securin-Ub550 binding event.
- If ZFAND5 signal appeared after Securin-Ub550 binding to proteasome but before the end of substrate processing or dissociation, this event was classified as “ZFAND5 binding during substrate on proteasome”.
- Finally, a ZFAND5 event was defined as “ZFAND5 binding after substrate dissociation” if the ZFAND5 signal appeared within 10 seconds after the Securin-Ub550 signal disappeared from proteasome.

#### Live-cell timelapse microscopy

HEK293T cells were transfected with fluorescent reporter constructs, Ub-R-YFP or UbG76V-YFP together with a plasmid expressing wildtype ZFAND5 or mutants from a CMV promoter using TransIT293 following the manufacture’s manual. 36 hours after transfection, cells were replated at about 40% confluency in a 12-well plate. After attachment, 100ug/ml cycloheximide or DMSO was added to the culture and cells were imaged in an Incucyte Zoom imager with 20X objective placed in a conventional tissue culture incubator every 7 minutes.

#### Calibration of the YFP reporter expression level

HEK293T cells were treated as above. 36 hours post transfection, cells were trypsinized. Cell density and cell volume were immediately measured using a TC20 cell counter. The total fluorescent intensity in the trypsinzed culture was determined using a BioTek H1 plate reader and compared with a purified YFP standard. The average cellular YFP concentration was calculated as [YFPculture]/(N*V), [YFPculture] is the equivalent YFP concentration in the trypsinzed culture; N: cell density; V: cell volume

#### Data analysis

Processing of timelapse images was performed using P53cinema which is an automatic cell tracking and segmentation software (Reyes et al., 2018). Briefly, each image was preprocessed, background subtracted. Individual cells were identified according to the local fluorescence maxima. The cell boundary was automatically segmented. Cells were automatically tracked along their movement. Tracking and segmentation results were manually verified. The software then calculated the average fluorescent intensity vs. time for each segmented cell. 100∼200 cells were analyzed for each condition. To extract the half live, each time trace was fitted with an exponential function to obtain the decay constant using MATLAB, and was registered with the initial reporter concentration calculated based on the calibration.

### QUANTIFICATION AND STATISTICAL ANALYSIS

Quantification of western blot and autoradiography results was performed in ImageJ. Quantification of timelapse images was performed using p53Cinema (Reyes et al., 2018). Quantification of single-molecule images was performed using custom software as described in a previous study (Lu et al., 2015). All statistical analysis was carried out in standard procedures using MATLAB 2021.

